# The landscape of antigen-specific T cells in human cancers

**DOI:** 10.1101/459842

**Authors:** Bo Li, Longchao Liu, Jian Zhang, Jiahui Chen, Jianfeng Ye, Alexander Filatenkov, Sachet Shukla, Jian Qiao, Xiaowei Zhan, Catherine Wu, Yang-Xin Fu

## Abstract

Antigen-specific T cells can be orchestrated to kill cancer cells in immunotherapies but the utilities of the TCR information have not been fully explored. Here, we leveraged previous efforts on tumor TCR repertoire, and developed a novel algorithm to characterize antigen-specific TCR clusters. Joint analysis with gene expression revealed novel regulators for T cell activation. Investigation of single-cell sequencing data revealed a novel subset of tissue-resident memory T cell population with elevated metabolic status. Integrative analysis of TCR clusters with HLA alleles and cancer genomics data identified candidate antigens derived from missense mutations, frameshift indels, and tumor-associated gene overexpression. Predicted antigen *HSFX1* was further validated using vaccinated humanized HLA-A*02:01 mice. Finally, high abundant cancer-associated TCRs were observed in the blood repertoire of early breast cancer patients, suggesting new avenues for noninvasive early detection. Thus, our analysis identified cancer-associated T cells with broad utilities in immune monitoring and cancer immunotherapies.

## Introduction

Antigen-specific tumor-infiltrating T lymphocytes (TIL) play a central role in cancer immunity^1-3^, with demonstrated applications in cancer immunotherapies, including checkpoint blockade^4-6^ and adoptive cell transfer therapies^7,8^. Therefore, identification of antigen-specific TIL is critical to understanding tumor-immune interactions and designing individualized treatments. However, this task remains challenging despite extensive clinical efforts^9,10^. First, cancer antigens may come from diverse sources, including missense mutations^11,12^, frameshift insertions or deletions^13,14^, tissue specific gene overexpression^15,16^, and other antigenic processes^17-20^, making it difficult to profile all the possible targets. In addition, the antigen-binding CDR3 region on the T cell receptor (TCR) is extremely diverse^21^, and their targets are usually unknown. Thus limited progress has been made in the analysis of TIL repertoire despite pressing clinical needs. This is because statistical significance is usually difficult to reach in such analysis unless using a large cancer cohort and a proper computational method, neither of which is currently available to study the tumor antigen-specific T cells.

Efforts have recently been made to partition the immune repertoire into groups linking to antigen-specificity (GLIPH)^22^, or evaluate the similarity of CDR3s with known specificity for functional predictions (TCRdist)^23^. However, TCRdist prediction relies on established antigen-binding TCR repertoire, which is usually unavailable for cancer studies, while GLIPH is benchmarked for infectious disease, and we demonstrated its suboptimal specificity to accommodate the extreme diversity of tumor antigens. Therefore, due to the complicated interactions between cancer and tumor-reactive TILs, more specific strategy is required to study the repertoire of cancer-associated TCRs for improved immune monitoring and immunotherapies.

In this work, we systematically identified the antigen-specific T cells using a novel CDR3 dataset profiled from over 9,700 tumor RNA-seq samples from the Cancer Genome Atlas (TCGA) and a new computational method for highly specific grouping of TCR CDR3 sequences. These unique resources allowed us to integratively analyze the antigen-specific TILs together with cancer genetic alterations, gene expression patterns, patient clinical profiles, single-cell RNA-seq data and immune repertoire sequencing data^24,25^ from the public domain. This pan-cancer analysis led to several interesting findings, which might not only provide novel targets for late stage cancer treatment, but also point to new avenues for preventive cancer screens. Specifically, our analysis identified a number of metabolic enzymes as potential negative regulators for T cell activation and revealed a novel tissue-resident memory T cell subpopulation for a subset of antigen-specific TILs. We also predicted candidate cancer antigens derived from somatic alterations or overexpressed cancer-associated genes with *in vivo* validations, which may expand the current pool of cancer antigens for future vaccination development. Finally, we developed a predictor from the antigen-specific CDR3s to distinguish cancer patients from healthy individuals, and demonstrated its potential application to non-invasive early cancer detection or immune monitoring.

## Results

### Detection of antigen-specific CDR3 clusters with iSMART

We have previously described the TRUST algorithm^26^ for sensitive detection of T cell receptor hypervariable CDR3 sequences using bulk tissue RNA-seq data. In this work, we applied a later version of TRUST^27^ with improved sensitivity to 9,709 TCGA tumor RNA-seq samples and assembled 1.5 million CDR3 sequences (**Figure 1a**). Of these, 170,000 were complete productive CDR3s, following the IMGT nomenclature^28^. A sizeable fraction of the human T cell repertoire is public, derived from biased V(D)J recombination^29^, and are present in both healthy and diseased individuals. To exclude the irrelevant public TCRs, we compared the TCGA TIL CDR3s with a large cohort of TCR repertoire data from non-cancer individuals^30^ (**Methods**). CDR3s observed in these samples with high abundances were excluded, leaving 82,000 non-public sequences.

**Figure 1.**
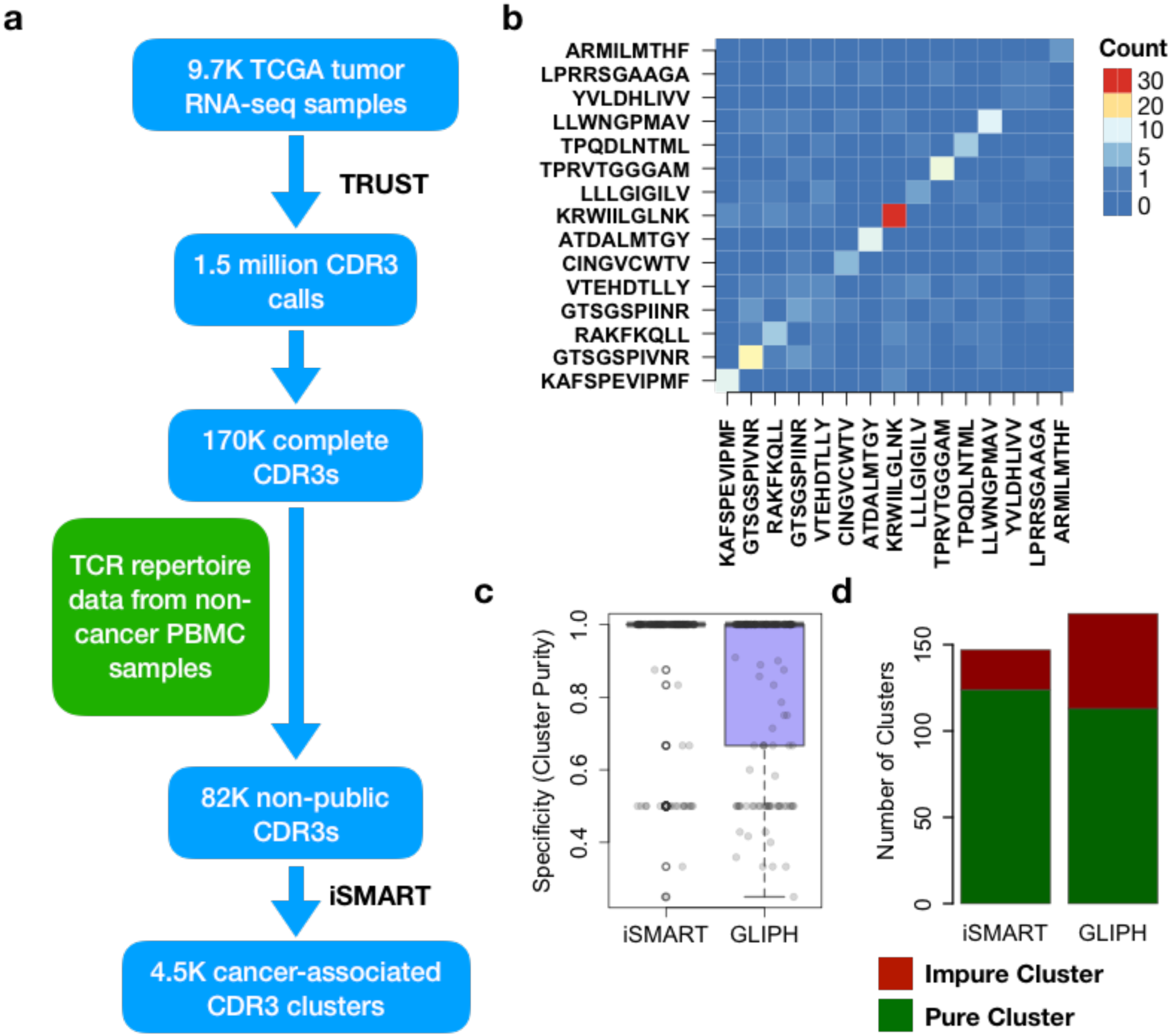
Methodology summary and performance evaluation for iSMART. **a)** Flowchart illustrating the analytical procedures carried out in this work to generate CDR3 clusters using TCR receptor sequences extracted from the tumor RNA-seq data. **b**) Heatmap representation for cross-antigen classification errors for iSMART. Each entry in the off-diagonal matrix is an integer representing the number of CDR3 groups showing co-clustering of the two antigens. Diagonal entries recorded the number of clusters with at least 2 CDR3s assigned to the corresponding antigen. **c**) Specificity comparison between iSMART and GLIPH. Cluster purities were displayed with boxplots, where 84.3% and 67.3% clusters have purity equal to 1 for iSMART and GLIPH respectively. **d**) Barplots showing the number of pure clusters for the two methods as a sensitivity measure to detect antigen-specific clusters.

Identification of antigen-specific CDR3 groups from TCR repertoire data is highly desirable, yet challenging due to the high diversity of CDR3 regions^21^ and promiscuous binding between T cell receptors and antigenic peptides^31-33^. A previous work, GLIPH^22^, demonstrated that CDR3s grouped into motif-sharing clusters are expected to recognize the same antigens. In our benchmark analysis we observed GLIPH groups a substantial fraction of CDR3s of different antigens (**Supplementary Figure 1**), thus might not be optimal for analyzing TIL TCR data. Therefore, we developed a new method, immuno-Similarity Measurement by Aligning Receptors of T cells, or iSMART, with increased clustering specificity (**Methods**). In brief, iSMART performs a specially parameterized local alignment on CDR3s, builds a pairwise comparison matrix and divides it into clusters with highly similar sequences. We benchmarked iSMART without variable (V) gene assignment, because 1) a large fraction of the TRUST assemblies do not have V gene information and 2) GLIPH does not rely on V gene assignment for clustering. We tested both methods using a curated CDR3 database containing the experimentally validated, antigen-specific TCR sequences^34^ (**Methods**).

We first applied iSMART to the 2,347 curated CDR3s specific to 15 selected antigens **(Supplementary Table 1**), and observed more specific grouping compared to GLIPH measured by cross-classification errors (**Figure 1b**). Overall GLIPH clustered a higher percentage of total CDR3s (29%) than iSMART (17%), but iSMART achieved significantly higher specificity (p=0.00078, Wilcoxon rank sum test) measured by cluster purity (**Figure 1c**) and called more clusters with unique antigen assignments (**Figure 1d**). As our goal is to identify tumor-specific CDR3s, the higher specificity of iSMART is a desirable feature. Therefore, we applied it to the 82,000 non-public CDR3 sequences and detected 4,501 clusters (**Figure 1a**). As most clusters contain more than one individuals, we also used the term ‘CDR3 cluster’ to denote the subset of patients carrying the CDR3s in a given cluster. A total of 15,254 CDR3 sequences were grouped into these clusters, and were referred as cancer-associated CDR3s.

### Features of CDR3 clusters and association with tumor gene expression profiles

The number of sequences in the clusters spans two orders of magnitude (**Supplementary Figure 2a**), and for each sample, the number of clustered CDR3s (***K***) also spans two orders of magnitude (**Supplementary Figure 2b**). For each gene, we calculated the partial Spearman’s correlation between ***K*** and its expression levels (**Supplementary Table 2**), controlled for tumor purity, which is expected to influence both values^35^ (**Methods**). Among the genes with top positive correlations are putative T cell activation markers, including *TBX21 (T-bet)*, *ICOS*, *TIGIT* and granzymes **(Supplementary Figure 3a**). Gene ontology enrichment^36^ analysis suggested that the top 500 genes are strongly enriched for immune cell activation and immune responses (**Supplementary Figure 3b**). Interestingly, on the top of the list there is a pair of lysophosphatidylserine receptors, *GPR174* and *P2RY10*, which have been identified as suppressors for regulatory T cell function^37^ (**Supplementary Figure 3a**). These results strongly suggest that CDR3s clustered by iSMART are enriched for activated T cells in the tumor microenvironment.

We next investigated genes negatively correlated with ***K*** as potential regulators for T cell inactivation and exclusion (**Supplementary Figure 4**). Of the 414 genes with correlation < -0.1 and FDR<0.05 in at least 3 cancer types, we observed 4 interesting clusters. Cluster i) contains a putative oncogene *MAPK3*^38^, inhibition of which has been linked to enhanced anti-tumor immune response^39^. This cluster also harbors a key glycolysis enzyme, *ALDOA*, which has recently been shown to impair T cell infiltration and cytotoxicity^40^. Cluster ii) contains two oncogenes, *RHOD* and *PKP3*, the former recently being implicated in immune suppression^41^. We also identified a number of other metabolic enzymes, including protein metabolic enzyme *POMGNT1*, cytochrome c enzymes *COX6A1* and *UQCRQ*, lipid metabolic enzymes *DGAT1* and *FAAH*, etc, supporting the recently elucidated immunosuppressive role of cancer metabolism pathways^42^.

### Identification of tissue-resident memory T subpopulations with distinct metabolic status

To further elucidate the phenotypes of the T cell clonotypes with clustered CDR3s, we analyzed a recently generated single cell RNA-seq (scRNA-seq) data with matched TCR information^43^. Using the TCGA-derived CDR3s as clonotype markers, we identified a number of clustered T cell clones in the 3 breast tumor scRNA-seq samples. We first studied sample BC10, which has the largest amount (n=55) of cells carrying clustered CDR3s. The selected 55 cells were visualized on the background of all 4,926 cells using t-Distributed Stochastic Neighbor Embedding (tSNE)^44^ plots, and observed a local clustering of 18 events in a restricted region (**Figure 2a**). All 18 events share the same β chain CDR3 sequence, and we delineated the region containing these cells as a separate CD8+ subgroup (n=44). Differential gene expression analysis on the cells in this group against all the others (**Supplementary Table 3**) revealed up-regulated genes both involved in T cell cytotoxicity (*GZMB*, *PRF1*, *IFNG*) and exhaustion (*PDCD1*, *LAG3*) (**Supplementary Figure 5**). Interestingly, the top targets showed high consistency with a recently reported tissue resident memory T cells (T_rm_) signature^45^, including up-regulation of *CD103* (*ITGAE*), *TIGIT*, *GZMB*, and down-regulation of *SELL (CD62L)*, *KLF2* and *KLRG1*. This group also expresses a number of other previously reported Trm markers^46^ (**Supplementary Figure 6**), including transcription factor *ZNF683*, or *HOBIT* (homolog of *Blimp-1* in T cells), a key regulator for T_rm_ differentiation^47^. We observed significant association of *ZNF683* expression with better outcomes in multiple cancer types (**Figure 2b**), supporting the anti-tumor role for T_rm_ cells.

**Figure 2.**
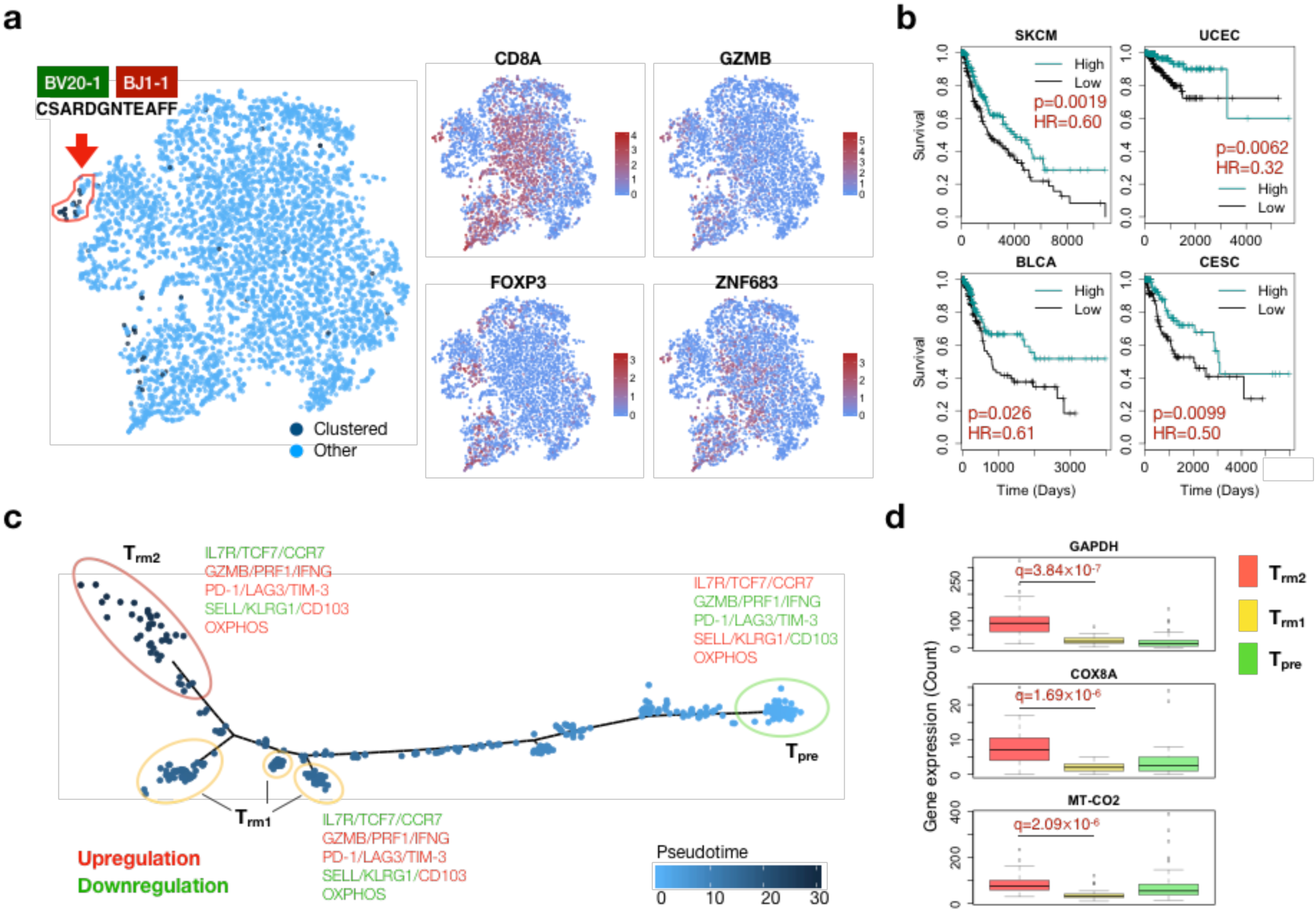
iSMART-clustered clonotypes showing tissue-resident memory phenotype. **a)** tSNE plots showing the distributions for clustered clonotypes in the TIL population (left), and the expression levels of selected putative markers for cell identity (*CD8A*/*FOXP3*/*ZNF683*) or function *(GZMB*) (right four panels). All selected markers passed FDR=0.05. Color legends for gene expression were in log scale. **b**) Kaplan-Meier curves for four TCGA cancers showing the survival benefit for *ZNF683* high expression. For each cancer, median value was applied to define high or low groups. Statistical significance and hazard ratio were evaluated using Cox proportional hazard model. **c**) Pseudotime trajectory plot illustrating the inferred evolutionary path. Cell clusters located on the beginning or end of the trajectory were manually selected. Representative markers significantly correlated (Spearman’s correlation test, FDR<0.05) with pseudotime inference were labeled for each cluster, with red for negative (high in T_pre_) and green for positive (high in T_rm_) correlations. **d**) Boxplots showing the distributions for selective OXPHOS genes in the three cell clusters shown in **c**). Statistical significance for differential gene expression between T_rm1_ and T_rm2_ was evaluated using Wilcoxon rank sum test, with FDR corrected by Benjamini-Hochberg method.

T cells undergo profound differentiations in the tumor microenvironment, and it is unclear which evolutionary path T cells have taken to become resident memory cells. The 44 cells in the subgroup come from 20 productive clonotypes, which in total contain 418 cells. We performed single cell trajectory analysis^48^ to infer the progression of these TILs (**Figure 2c** and **Methods**). The pseudotime trajectory starts from a group of precursor cells (T_pre_) expressing high levels of *IL7R*, *SELL*, and *KLRG1*, with low expression of effector molecules (*GZMB*, *PRF1*, *IFNG*) and exhaustion markers (*PDCD1*, *LAG3, TIM-3*). These markers agree with the signatures of T cells newly entering the tumor microenvironment, thus confirming the pseudo temporal ordering. Two clusters were observed at the end of the trajectory, both carrying the resident memory markers, and we named them T_rm1_ and T_rm2_. Notably, the T_rm1_ cluster largely overlaps with the previously identified CD8+ subgroup. Differential expression analysis revealed that comparing to T_rm1_, the newly identified T_rm2_ population upregulates genes significantly enriched in the oxidative phosphorylation (OXPHOS) process (FDR=6.77×10^-33^), featured by *GAPDH*, *COX8A* and *MT-CO2* (**Figure 2d**). Pseudotime trajectories for individual clonotypes revealed that the differentiation of T cells into resident memory status is receptor dependent (**Supplementary Figure 7**). Specifically, we observed two modes of evolution: majority of clonotypes evolve from T_pre_ into T_rm1_, with others from T_rm1_ to T_rm2_. Direct differentiation of T_pre_ into T_rm2_ was not observed.

We analyzed other scRNA-seq samples to see if this observation is reproducible, and indeed, a strikingly similar pseudotime trajectory for resident memory T cells was observed in sample BC11 **(Supplementary Figure 8a**). Representative markers observed in sample BC10 also showed significant differences across the 3 cell groups with consistent trends. Higher expression of OXPHOS genes was also observed in T_rm2_ group (**Supplementary Figure 8b**). In addition, the corresponding clonotypes also displayed two evolutionary patterns (**Supplementary Figure 8c**), consistent with our findings for BC10. These results indicate that the T_rm_ cells further divide into two populations distinguished by low or high metabolic status, and the differentiation from their precursors into these populations is dependent on the T cell receptors.

### Identification of novel cancer neoantigen candidates

Computational identification of cancer neoantigens relies on the prediction of peptide binding to HLA alleles^49,50^, while little is known whether the predicted binders can elicit an immune response. Having studied the phenotypes of the clustered T cell clonotypes using single-cell sequencing data, we next sought to approach this problem from the T cell receptor angle using the iSMART-derived CDR3 clusters and TCGA cancer genomics data. We have previously provided a proof-of-concept analysis to statistically identify novel candidate neoantigens^26^. In this work, we extended this effort by searching for co-occurrence of somatic mutations and CDR3 clusters (**Methods**). First, each of the 6,136 recurrent (n≥3) missense mutations was paired with each of the CDR3 clusters, with statistical significance evaluated using random permutations. At FDR<0.05, we identified 6 significant pairs with at least 2 individuals carrying both the mutation and the CDR3 sequence **(Figure 3a**). 4 of them (excluding the two *CD163L* mutations) are predicted to generate HLA binding peptides. Individuals carrying two of these mutations, *GLIS3* S47L and *SLITRK3* E968K, have matched HLA genotypes (**Figure 3b** and **Supplementary Table 4**).

**Figure 3.**
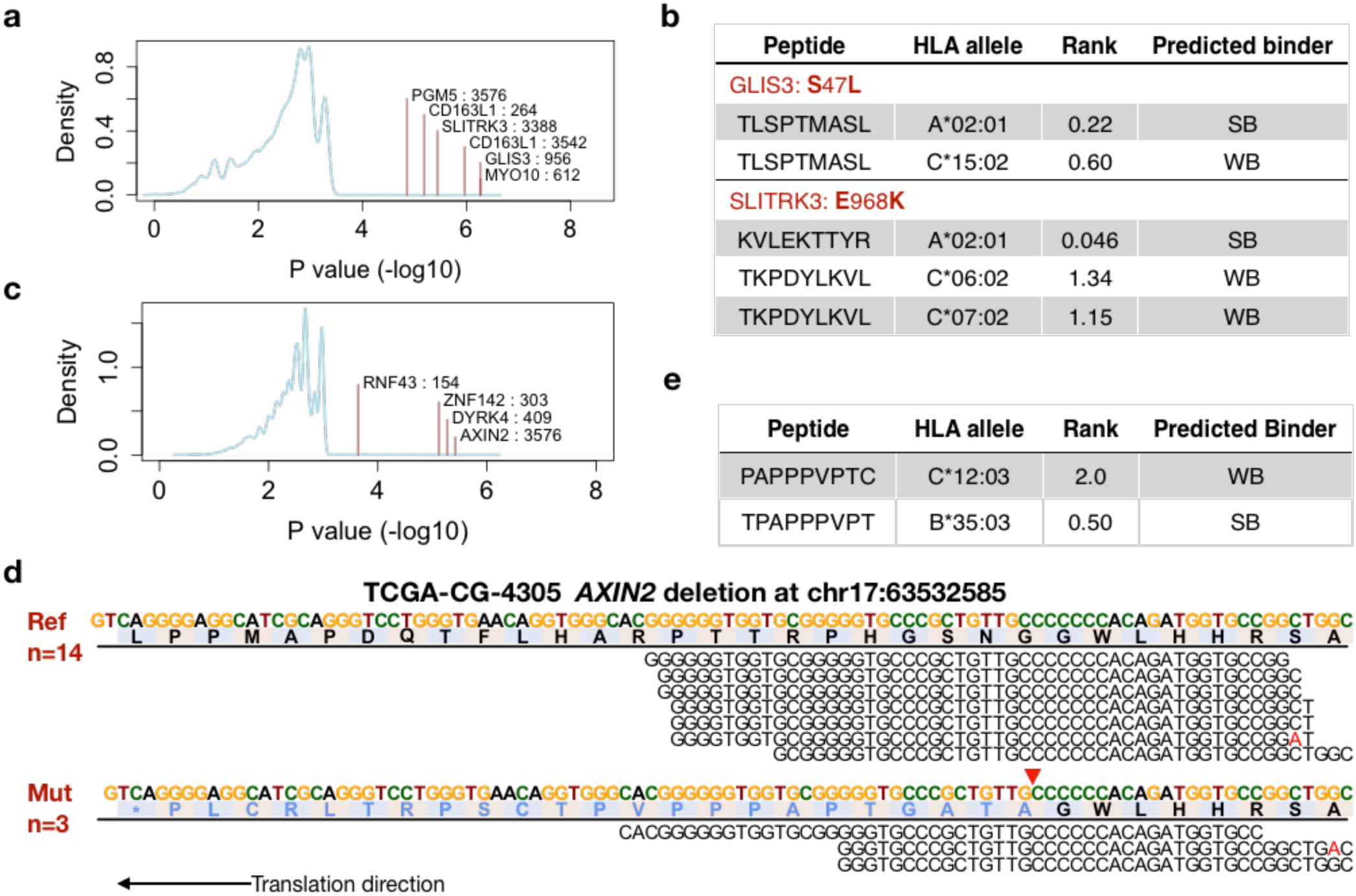
Prediction of neoantigens derived from missense somatic mutations and frameshift indels. **a**) Probability density plot for the null distribution of p values testing the significance of co-occurrence between CDR3 clusters and recurrent missense mutations. Significant mutations were marked in the plot with vertical lines and text labels. The numbers after each gene symbol are CDR3 cluster IDs. **b**) Table showing the NetMHC predicted binder peptide sequences, related HLA alleles and binding strength classification. **c**) Density plot for the p values from the analysis for frameshift indels. **d**) Reads pileup plot for reference and mutated alleles in the site with frameshift deletion in gene *AXIN2*. Numbers of total reads for either reference or mutated alleles were presented in the text box on the left. The gene is reversely translated and a stop codon (*) is generated 23 amino acids downstream of the site of deletion (red arrow). Mismatches were labeled with red color. **e**) NetMHC predictions for peptide binders generated from the frameshift deletion of AXIN2.

It has been implicated that tumor frameshift insertions or deletions (indels) may generate neopeptides to trigger immune responses^14^. We applied the same analysis to study the 1,225 recurrent (n≥3) frameshift indels. Compared with missense mutations, we observed more significant pairs (n=10) with fewer indels, indicating that frameshift indels might be another important source of neoantigens. As aberrant mRNA products are subject to nonsense-mediated decay^51^, we only focused on these (n=4) with frameshift alleles confirmed in the RNA-seq data **(Figure 3c**). The top target is a one-base deletion in gene *AXIN2*, generating a 23 amino acids neopeptide (**Figure 3d**). We predicted HLA binding for the neopeptide, and identified two closely related 9-mers to bind to the HLA alleles of the mutation carriers (**Figure 3e**). Besides *AXIN2*, two other indels on *DYRK4* and *RNF43* also generate neopeptides predicted to bind to HLA alleles matching the genotypes of the carriers (**Supplementary Figure 9**). We observed that all the related individuals were stomach cancer patients with high levels of microsatellite instability (MSI)^52^, and the indels all occurred in the short tandem repeat regions. It is known that patients with DNA mismatch repair (MMR) deficiency, a cause for MSI, have better responses to checkpoint blockade therapies^53^. Our results corroborate this clinical observation by identifying a number of potentially immunogenic neoantigens resulted from MMR deficiency, and demonstrated that iSMART can prioritize TCRs associated with neoantigens derived from genetic alterations.

### Identification of novel cancer associated antigen candidates

Most current studies focus on searching for tumor antigens from mutated genes with matched HLA alleleotypes combining the elution of peptides from the MHC molecules^54^. However, malignant cells may overexpress a number of genes that are usually silenced in most normal tissues, resulting in novel antigenic targets for cancer treatment. This is exemplified by the clinical use of cancer/testis antigens, which have restrictive expression in the male germ cells^15,16^. We performed a genome-wide differential gene expression analysis on each of the 120 qualifying CDR3 clusters, and identified a total of 1,409 significant (FDR<0.05) genes from 115 clusters **(Methods** and **Supplementary Table 5**). Of these, two clusters (1724 and 1767) showed an interesting enrichment in colon and endometrial cancers, with distinct CDR3 conservation patterns (**Figure 4a**). We performed differential expression analysis on the combined samples from the two clusters, and identified Heat Shock Transcription Factor X-linked 1 (*HSFX1*) as the top hit (**Figure 4b**). This gene has extremely low expression (median TPM≤0.02) in all the tissue types covered in the GTEx data^55^, while expressed (TPM≥1) in 13% colorectal and 75% endometrial cancers (**Supplementary Figure 10a-b**). There is over 100-fold change in the expression levels between some tumor samples and the normal tissues. It is also a favorable predictor of survival for endometrial cancer (**Figure 4c**). Therefore, we hypothesized that the tissue-specific overexpression for *HSFX1* may be a signature for cancer-associated antigen and a trigger for anti-tumor immune response.

**Figure 4.**
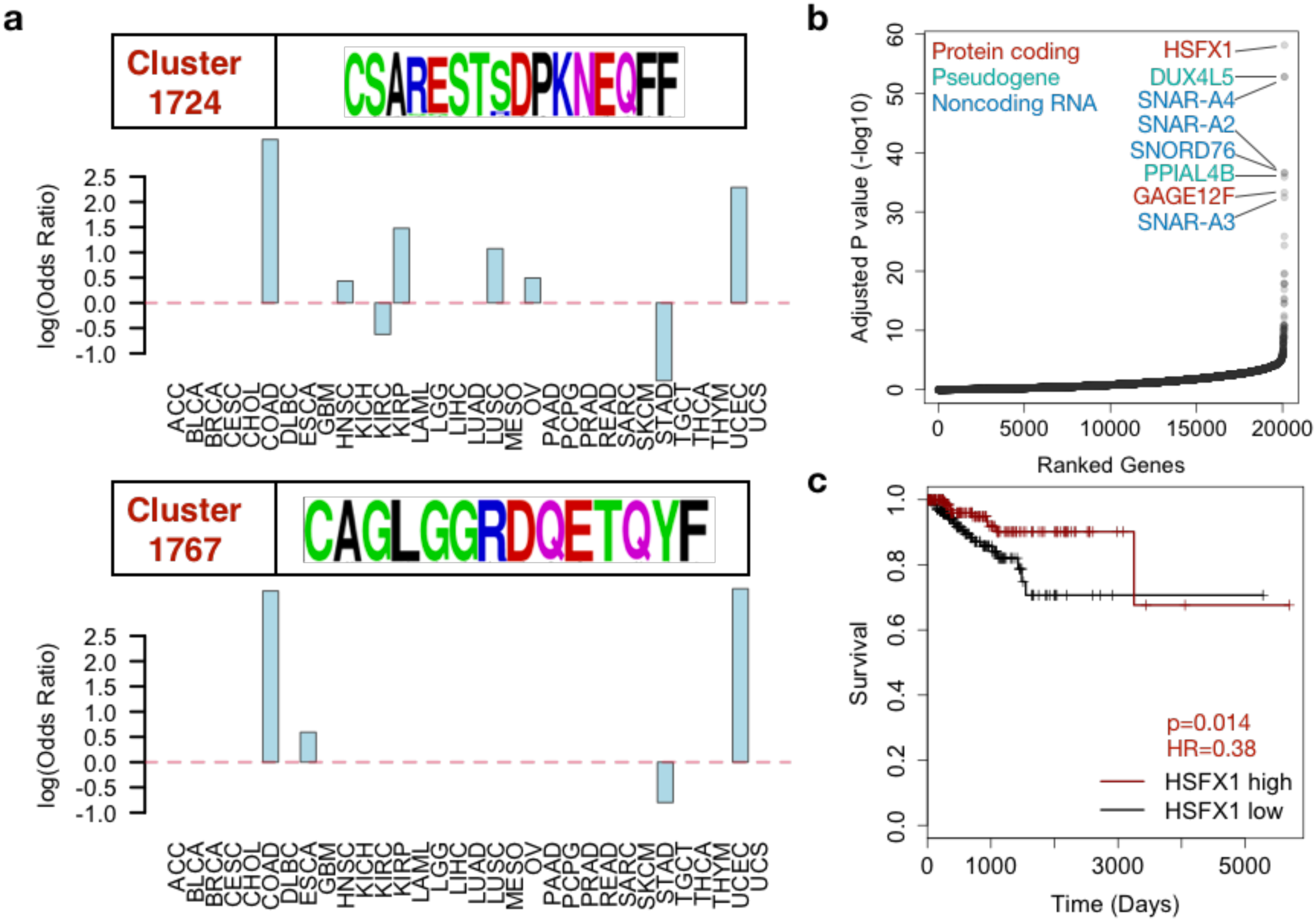
Identification of HSFX1 as a candidate cancer-associated antigen. **a**) Selective enrichments in colon and endometrial cancers of samples in CDR3 clusters 1724 and 1767. CDR3 amino acid conservation patterns were displayed in the upper panel for each barplot. **b**) Genes ranked by p values from differential gene expression analysis, with top hits labeled in colored texts. HSFX1 has the most significant p value among all the genes. Statistical significance was evaluated using Wilcoxon rank sum test with FDR correction. **c**) Kaplan-Meier survival curves for endometrial cancer patients with or without HSFX1 expression, separated by median expression value. Statistical significance and hazard ratios for HSFX1 levels were estimated using Cox proportional hazard model on binary input of HSFX1 groups, corrected for patient age.

Of the seventeen colon or endometrial cancer samples from cluster 1724 and 1767, nine express *HSFX1* and have solved HLA genotypes^56^ (**Supplementary Figure 11a**). Computational prediction for HLA allele binding suggests that *HSFX1* protein generates a 9-mer peptide VMFPHLPAL as a strong binder to 3 common alleles, including HLA-A*02:01 (**Supplementary Figure 11b**). Interestingly, all the nine individuals carry at least one predicted HLA allele (**Supplementary Figure 11c**), and the probability of this observation is estimated ≤0.00038 (**Methods**). These results strongly suggested that *HSFX1* might be an immunogenic cancer antigen. To validate this prediction, we synthesized the 9-mer antigen peptide (VMF) and injected it into HLA-A*02:01 humanized transgenic mice (**Methods**). We used peptide V**R**FPHLPAL, which has one amino acid difference, as control, because it is predicted not to bind HLA-A*02:01. After 18 days, splenocytes of the vaccinated mice were collected to perform an IFNγ ELISPOT assay for antigen-specific T cell responses (**Figure 5a**). Compared to the control peptide (VRF), we observed significantly higher IFNγ response in the transgenic mice but not in identically primed immunocompetent mice with H-2K^b^ genotype (**Figure 5b-c, Supplementary Figure 12**). Based on these results, we concluded that the 9-mer peptide VMF derived from *HSFX1* binds to human HLA-A*02:01 allele and can elicit a T cell response *in vivo*.

**Figure 5.**
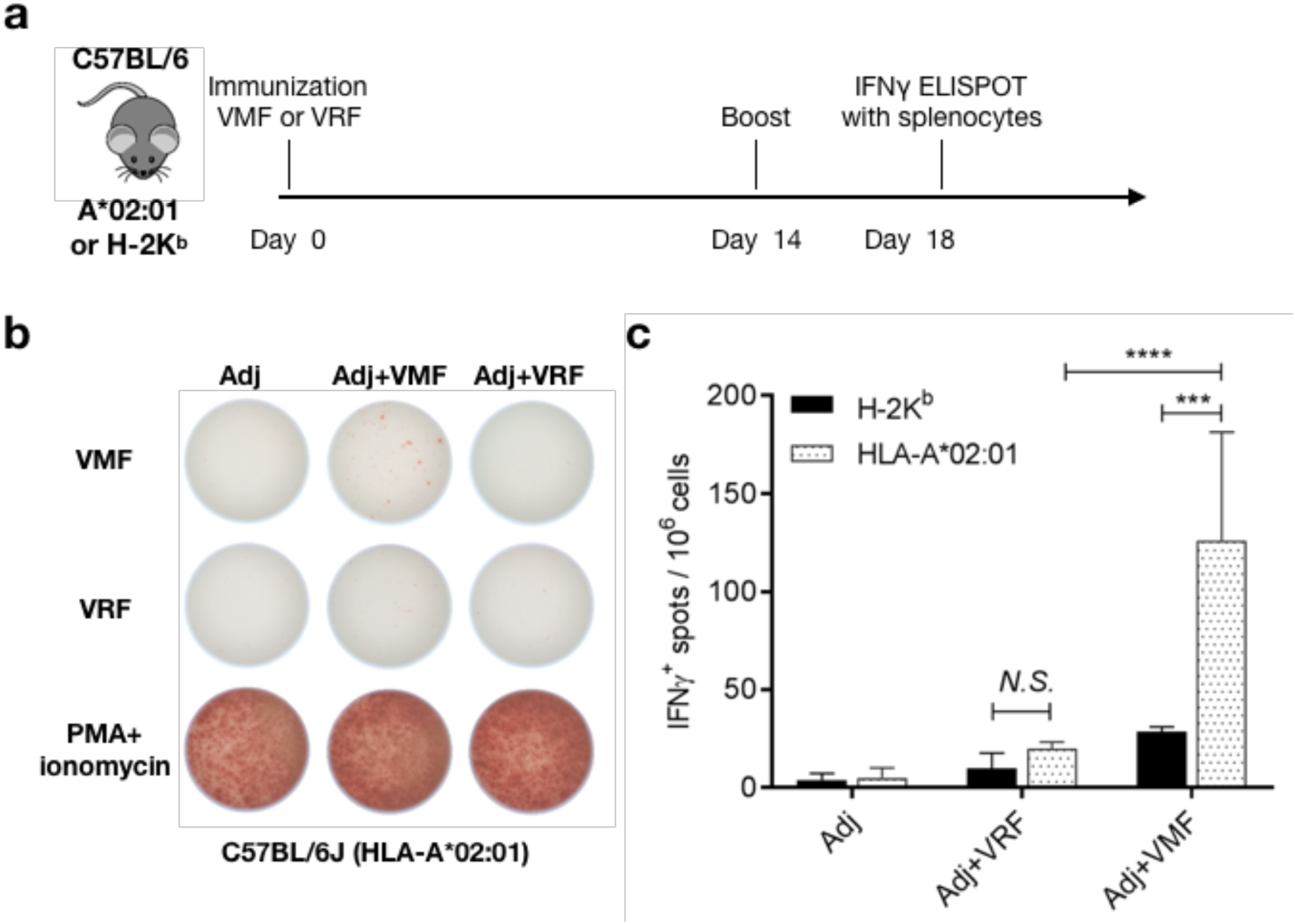
Immunogenicity of the 9-mer peptide derived from HSFX1 protein in HLA-A*02:01 transgenic mice. **a)** HLA-A*02:01 transgenic mice (female, n=4) were subcutaneously immunized with 10μg peptide mixed with 100μg poly(I:C) and 50μg CpG1826. 14 days post vaccination, mice were boosted with the same vaccine. 4 days later, splenocytes were isolated for IFNγ ELISPOT assay. Representative results showed IFNγ secreting cells from indicated groups (**b**). Column texts labeled the 3 treatment groups, where the T cells were collected. Row texts labeled simulants used in the ELISPOT assay. Significant difference of antigen-specific T cell response from the control peptide was observed (**c**). Data are expressed as the means ± SD, representative results from two independent experiments are shown. Statistics analysis was performed by Two-way ANOVA. ***, P < 0.001 ; ****, P < 0.0001. Adj is short for adjuvant injected during vaccination. VMF: antigen peptide VMFPHLPAL; VRF: control peptide V**R**FPHLPAL; PMA+ionomycin: standard positive control.

In addition to *HSFX1*, we also identified a putative cancer/testis antigen, *TSSK2*, with expression restricted to esophageal and stomach tissues (**Supplementary Figure 13a**). *TSSK2* also generates a peptide binding to multiple HLA alleles (**Supplementary Figure 13b**), matching the genotypes of the individuals expressing *TSSK2* from cluster 189 (**Supplementary Figure 13c),** with probability <0.0010 (**Methods**). These results suggest that genes with tumor-specific overexpression might produce cancer-associated antigens and elicit T cell responses. Our analysis revealed a number of such unmutated genes as promising targets for cancer vaccine development.

### Potential non-invasive cancer diagnosis using cancer-associated CDR3s

In the above analysis, we observed multiple sources of potential tumor antigens showing significant associations to the iSMART identified CDR3 clusters, suggesting that the clustered CDR3s are enriched for cancer-associated T cells. We therefore investigated whether it is possible to detect these CDR3s in the TCR repertoire from the peripheral blood mononuclear cell (PBMC) of the cancer patients. We studied a cohort of 21 late-stage melanoma patients before and after anti-CTLA4 treatment^25^. When compared to the healthy donors, we identified significantly higher abundance of cancer-associated CDR3s in the patients’ blood samples (**Figure 6a**) (**Methods**). Using cancer-associated CDR3 counts as a disease predictor, pre- and post- PBMC samples reached similar area under curve (AUC) of 0.80 and 0.82 respectively (**Figure 6b**).

**Figure 6.**
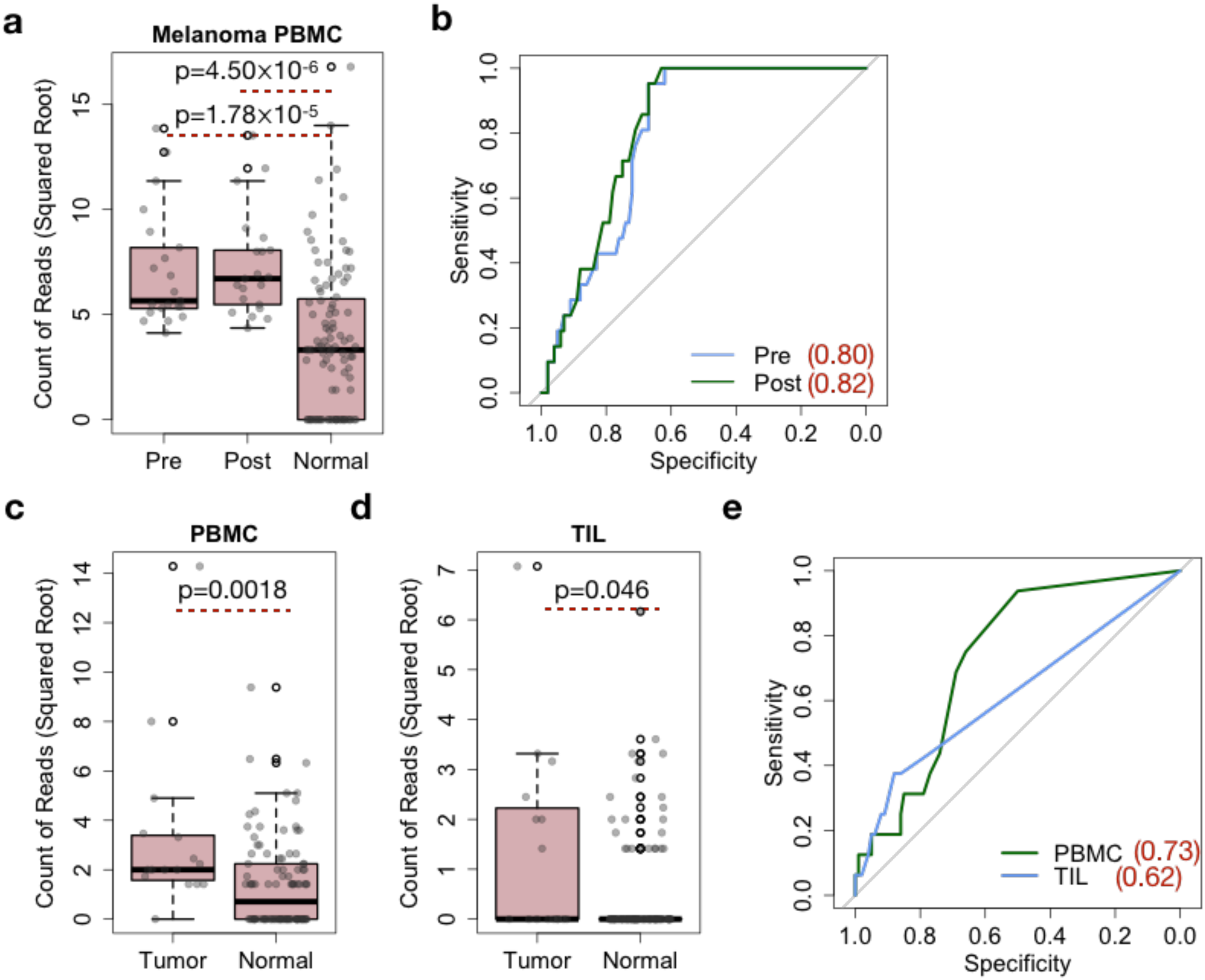
Prediction of late and early stage cancers using cancer-associated CDR3s. **a**) Boxplot showing the distributions of the read counts for cancer-associated CDR3s for pre- or post-anti-CTLA4 treatment late-stage melanoma and normal control samples. TCR repertoire data from all the samples were derived from PBMCs. **b**) ROC curves for using CDR3 read count as a predictor for late stage melanoma. Numbers in the figure legend are area under curve (AUC) values. **c-d**) Cancer-associated CDR3 read count distributions for early stage breast cancers comparing to normal samples, with cancer samples being PBMC (**c**) or TIL (**d**). **e**) ROC curves for using the abundance of cancer-associated CDR3s PBMC or TIL samples as predictors for early breast cancer onset. AUC values were shown in the legend. Statistical significance was evaluated using Wilcoxon rank sum test between labeled groups.

We next evaluated the performance of the above approach on the challenging yet more useful task of predicting early cancer status via PBMC repertoire. We applied the same method to study a cohort of 16 early-stage breast cancer samples with both PBMC and TIL repertoires sequenced^24^. Indeed, both repertoires showed significantly higher levels of cancer-associated CDR3s than healthy donors (**Figure 6c-d**), indicating that the abundance of cancer-associated CDR3s is able to distinguish healthy individuals from both late and early stage cancer. Using iSMART-clustered CDR3 counts as a predictor, we observed an AUC of 0.73 for PBMC samples (**Figure 6e**). With more future studies on pre-cancer TCR repertoire sequencing, this approach holds the potential to be developed into a non-invasive cancer diagnostic criterion.

The distributions of the clonal frequencies of the cancer-associated CDR3s in the PBMC samples also showed interesting differences between late and early stage tumors (**Supplementary Figure 14**). Specifically, melanoma PBMC samples have more cancer-associated CDR3s with medium high abundance, where early breast cancer PBMC samples have a few CDR3s with high abundance. This might be related to the known fact that in an adaptive immune response, many effector T cells differentiate into memory cells for long-term protection, resulting in reduced clonotype frequencies. Based on a previous study on cancer and inflammation^57^, we speculated that during early cancer development, the immune system is able to recognize and respond to a few shared antigens (such as *HSFX1*), and produce a significant amount of effector T cells in the circulation. The CDR3 sequences of these T cells may serve as diagnostic markers for preventive early cancer detection or immune monitoring.

## Discussion

Despite extensive efforts and critical clinical applications, antigen-specific TILs remain largely uncharacterized, mainly because it is experimentally challenging to identify the immunogenic cancer antigens and to profile the tumor-reactive T cells. In this work, we extracted CDR3s from the tumor RNA-seq data, and identified a large number of CDR3 clusters with high sequence similarity. Due to the excessive diversity of the TCR repertoire, the probability that different individuals independently produce near-identical non-public TCRs is extremely low, suggesting that shared antigen-specificity is the main cause for the generation of these CDR3 clusters. Previous studies have also shown that TCRs sharing motifs on the CDR3 region may recognize the same antigen^22,23^. Therefore, we used iSMART identified CDR3 clusters as surrogates for TCR antigen-specificity, and comprehensively analyzed the tumor-specific TILs using a large human cancer cohort.

We leveraged the iSMART-clustered clonotypes to perform an in-depth analysis of a tumor single cell RNA-seq dataset with solved T cell receptor sequences, and observed an interesting group of CD8+ T cells. The marker set for this group is highly consistent with a recent study on T_rm_^45^, suggesting reproducible identification of T_rm_ in triple-negative breast tumor microenvironment. Using CDR3 as clonotype markers, we further identified two subpopulations of T_rm_ with distinct metabolic states, and observed divergent evolutionary paths to these states among different TIL clonotypes. Our results suggest that after initial homing to the target tissue, T_rm_ may switch to a high metabolic status, featured by elevated expression of OXPHOS genes. This result is potentially linked to the immunosuppressive roles for metabolic enzymes in the malignant cells, which they use to compete resources for T cell survival and cytotoxic functions.

It has been shown from protein structure studies that one antigenic peptide may bind to dissimilar CDR3 sequences with different docking strategies^58,59^, suggesting that individuals responding to the same antigen may carry divergent TCR sequences. Indeed, we observed two distinct CDR3 sequences from clusters 1724 and 1767, which were both predicted to recognize the same antigen derived from cancer-associated antigen *HSFX1*. We performed *in vivo* experiments using transgenic humanized mice to show that a 9-mer peptide derived from a predicted antigen *HSFX1* is able to bind HLA-A*02:01, and induce reliable T cell responses. These results, combined with the observation that *HSFX1* has restricted expression in selected cancers, and its positive clinical relevance, strongly indicated that it might escape central tolerance in humans and become an immunogenic cancer-associated antigen. We will rely on future clinical studies using colorectal or endometrial cancer patients expressing *HSFX1* to explore its potential clinical utilities.

A fraction of the CDR3 clusters remain unassociated with any potential targets, likely due to the unexplored categories of cancer-associated antigens. In our gene expression analysis, we observed significant associations of some clusters with non-coding RNAs (**Supplementary Table 5**), such as lncRNA, pseudogenes and small nucleolar RNAs (snoRNA). Ribosome profiling data suggests that many non-coding RNAs are actually translated^60^, which may serve as valid cancer antigens when overexpressed in the tumor tissues. snoRNAs participate in many biological processes, including RNA splicing. Thus, their abnormal expression in selected cancer types may produce new antigenic targets from alternative splicing. Post-translational modification (PTM) may also generate foreign peptide products that are subject to immunosurveillance^17^. However, due to the insufficiency of related data, it is currently challenging to study the antigenic potentials of these mechanisms in cancer immunity.

The iSMART identified CDR3 clusters might have promising applications in cancer diagnosis. In our proof-of-principle analysis on a small patient cohort, we observed promising predictive power for early-stage breast cancers using blood TCR repertoire data. This result is reproducible for late stage melanoma, suggesting that the clonal expansion of cancer-associated CDR3s in the PBMCs might be universal to cancer types and stages. Therefore, we anticipate more clinical efforts to collect PBMC repertoires profiled from early stage cancer patients to elucidate the feasibility of this non-invasive approach for cancer detection, or immune monitoring during cancer therapies.

In summary, we provided a comprehensive analysis to characterize cancer antigens and tumor-reactive T cells. The tool and datasets from this study can be applied to the rapidly generated tumor single-cell sequencing and RNA-seq data to expand the current repertoire of cancer-associated TCRs. Therefore, we anticipate broad utilities of our work for future studies to identify more antigens and biomarkers for cancer immunotherapies.

## Abbreviations

TCGA: The Cancer Genome Atlas
CDR3: Complementarity Determining Region 3
ACC: adenocortical carcinoma
BLCA: bladder carcinoma
BRCA: breast carcinoma
CHOL: cholangiocarcinoma
CESC: cervical squamous carcinoma
COAD: colon adenocarcinoma
DLBC: diffusive large B-cell lymphoma
ESCA: esophageal carcinoma
GBM: glioblastoma multiforme
HNSC: head and neck carcinoma
KICH: kidney chromophobe
KIRC: kidney renal clear cell carcinoma
KIRP: kidney renal papillary cell carcinoma
LAML: acute myeloid leukemia
LGG: lower grade glioma
LIHC: liver hepatocellular carcinoma
LUAD: lung adenocarcinoma
LUSC: lung squamous carcinoma
MESO: mesothelioma
OV: ovarian serous cystadenocarcinoma
PCPG: pheochromocytoma and paraganglioma
PAAD: pancreatic adenocarcinoma
PRAD: prostate adenocarcinoma
READ: rectum adenocarcinoma
SARC: sarcoma
SKCM: skin cutaneous melanoma
STAD: stomach adenocarcinoma
TGCT: testicular germ cell tumor
THCA: thyroid carcinoma
THYM: thymoma
UCEC: uterine corpus endometrial carcinoma
UCS: uterine carsinosarcoma

## Acknowledgement

This work is supported by CPRIT grant 50C1284401 (BL), Circle of Friends Cancer Center Grant 2018 (BL), and CPRIT grants RR150072 (YXF). We thank Dr. James Brugarolas for helpful discussions during manuscript preparation.

## Author contributions

BL conceived the project, developed the algorithm and performed the computational analysis. LCL designed and performed the in vivo experiment. JZ contributed to algorithm coding. JHC, JFY, XWZ, JQ and AF contributed to data analysis. SS and CW contributed data. BL and LCL wrote the manuscript. BL and YXF supervised the study.

## Data and code availability

iSMART source code is available at https://bitbucket.org/lilab_utsw/ismart/. The CDR3 clustering data is hosted on Firecloud (https://software.broadinstitute.org/firecloud/) and is available upon request under TCGA controlled access.

## Methods

### Data resources information

TCGA level-2 RNA-seq data aligned to hg19 human reference genome by MapSplice^61^ were downloaded from GDC legacy archive (https://portal.gdc.cancer.gov/legacy-archive/). Gene expression data (TPM), mutation annotation files and patient clinical information of the TCGA cohort were downloaded from GDAC broad firehose (https://gdac.broadinstitute.org/). Tumor purity information was downloaded from the Cistrome TIMER website (http://cistrome.org/TIMER/misc/AGPall.zip). TCR repertoire data and patient information for the HCMV cohort, late stage melanoma and early breast cancer samples, were downloaded from AdaptiveBiotechnology immunoSeq Analyzer (https://www.adaptivebiotech.com/). Antigen-specific CDR3 sequence information for benchmarking iSMART were downloaded from VDJdb (https://vdjdb.cdr3.net/). GLIPH software package was accessed from GitHub (https://github.com/immunoengineer/gliph). Single cell gene expression data and matched TCR information were downloaded from GEO database (accession number GSE114724).

### Materials and animal model

HSFX1 derived 9-mer peptide was synthesized by GenScript; CpG oligonucleotide ODN 1826 was purchased from InvivoGen with catalog number 1826-1; Polyinosinic-polycytidylic acid sodium salt, or Poly (I:C), was ordered from MiliporeSigma with catalog number P1530-25MG. Immunocompetent C57BL/6J and transgenic C57BL/6-Mcph1Tg(HLA-A2.1)1Enge/J mice were obtained from Jackson Laboratory (JAX:000664 and JAX:003475)

### iSMART for pairwise CDR3 alignment and clustering

iSMART takes *M* complete CDR3 sequences as input, where complete CDR3 region is defined as the last cysteine in the variable gene to the first amino acid in the FGXG motif in the joining gene^28^. iSMART first orders the CDR3s according to their lengths, and then performs pairwise comparisons for every sequence. For CDR3s with different lengths, iSMART allows at most one insertion in the comparison, and imposes a gap penalty (default 6). Alignment scores are calculated based on BLOSUM62 matrix, with individual matched score capped at 4. The 3^rd^ to (*n*-3)^th^ positions of the CDR3s are used for scoring, where *n* is the CDR3 amino acid sequence length. Pairwise alignment score is normalized by the length of the longer CDR3 sequence (*n*-4, excluding first and last 2 amino acids). After calculation of the *M*-by*-M* pairwise scoring matrix, a predefined cutoff value (default 3.5) is applied to filter out all the low scoring comparisons. iSMART then performs a depth-first search on the matrix to identify all the connected CDR3 clusters, and output all the CDR3s with empirical cluster IDs. iSMART is written in Python and the source code is publicly available.

Although iSMART is benchmarked to run without variable gene assignment in this work, it supports the input with variable gene information. In this mode, the pairwise alignment on the CDR3 regions is the same except that iSMART uses the 5^th^ to (*n*-3)^th^ positions of the sequence for scoring. As the first 4 amino acids of the CDR3s are mainly determined by the variable gene, we made this change to avoid repeated use of variable gene information. In the pairwise sequence comparison step, the CDR1 and CDR2 regions of two TCRs are also used to calculate alignment scores under the same rules. The total score is scaled to 8, where CDR3 and variable gene contribute equally, and a cutoff value (default 7.5) is used to generate the CDR3 clusters. iSMART in variable gene mode was tested using the 15 antigen benchmark dataset, which is described in section below, and reached a higher specificity of 94.3% (100 out of 106 clusters have unique antigen assignment) than without variable gene input.

### iSMART and GLIPH performance evaluation

Both iSMART and GLIPH can predict antigen-specific CDR3 clusters without variable gene information. In this work, we evaluated the performances of both methods using TCRs of known antigen-specificity in VDJdb^34^. We selected 15 9-mer human antigens with balanced number (K) of associated TCRβ CDR3s (100<K<1000) (**Supplementary Table 1**). CDR3s associated with more than one antigens were excluded, resulting in a total of 2,347 unique sequences. Both iSMART and GLIPH were run on this dataset with default parameters.

The command line for iSMART is:

~~~
python iSMARTv1.py –f human15aa.txt –v
~~~

where –v option is applied to disable the use of variable gene. For GLIPH the command line is:

~~~
./gliph-group-discovery.pl ––tcr human15aa.txt
~~~

Interestingly, although iSMART performs time-consuming pairwise sequence alignments, its computational time (63s) is significantly less than GLIPH (approximately 1 hour) on MacBook Pro with 3.1 GHz Intel core i7 and 16 GB DDR3 memory. Therefore, iSMART has the computational efficiency to scale up for larger TCR repertoire datasets.

As each CDR3 is uniquely linked to one antigen in the benchmark dataset, we defined cluster purity (*p*) as the number of the most abundant antigen divided by the number of CDR3s in a cluster. We use the percent of completely pure (*p*=1) clusters as a measure for specificity. To make visualization of the clustering specificity, we computed the cross-antigen classification errors as follows: the 15-by-15 cross-antigen matrix (*M*) is initialized by 0, and for each cluster, let A denote the set of antigens associated with the CDR3s in this cluster, we add 1 to all the entries in *M*[A, A]. Therefore, if A contains only one antigen, the diagonal values for *M* will increase by 1. Otherwise the off-diagonal values will increase by 1, which are considered as classification errors. We looped through all the clusters and used the final output to plot the heatmaps in **Figure 1** (iSMART) and **Supplementary Figure 1** (GLIPH).

### Non-cancerous public TCR identification

A critical pre-processing procedure in our analysis is to exclude non-cancerous public TCRs to reduce false positives in our downstream analysis. We used a cohort of non-cancer individuals with TCR repertoire sequencing data available^30^. There are two batches of this cohort, with the first batch containing 666 human cytomegalovirus (HCMV) infected (n=289) or normal individuals. The HCMV infected individuals can be used as control samples for our purposes. The second batch contains 120 individuals. We will use the first batch to remove public TCRs and the second for downstream analysis, to avoid systematic bias. Based on antigen-specificity, we processed two classes of public TCR sequences:

The first class is antigen-specific non-cancerous CDR3s. In our downstream analysis of detection cancer-associated CDR3s in the blood TCR repertoire, we also rely on this HCMV cohort as normal control. Therefore, at this step, to prevent any potential confounders, we used the first batch (n=666) to remove public TCRs. It is known that cancer-specific T cells are also present in healthy individuals in the form of low abundant naïve T cells^62^. Therefore, to prevent false removal of *bona fide* cancer-specific CDR3s, we restricted our analysis within the top 5,000 most abundant sequences, sufficient to cover all the clones with ≥5 copies that are expected to be effector T cells. We combined all the sequences as normal CDR3s to be removed in the TCGA data before iSMART clustering. The resulting dataset as well as samples used in this analysis are available as **Supplementary Dataset 1**.

The second class of public sequences are non-antigen specific CDR3s, potentially due to biased V(D)J recombination. We anticipate that the sharing of these sequences between individuals is not affected by the HLA alleles of the carriers. Therefore, we performed 800,000 random sampling of triplets from the pool of 666 TCR repertoire samples satisfying the following criterion: there is no overlap in the HLA alleles in any two individuals in the triplet. For each triplet, we compared the top 5,000 most abundant sequences in each sample and selected those appeared in all three. The resulting 3,470 CDR3 sequences are available as **Supplementary Dataset 2**.

We removed both classes of public sequences from the 170,516 complete CDR3s and obtained 82,427 non-public sequences for downstream analysis. As the TCR repertoire data in the public domain are mainly β chain sequences, currently we do not have enough data to eliminate public α chain CDR3s from the analysis. We will rely on future efforts to sequence more TCR α chain repertoire samples to define public α chain CDR3 sequences.

### Analysis of single cell sequencing data

Post-processed gene expression data in sparse matrix format (mtx) and TCR hypervariable CDR3 sequences with matched cell barcodes were downloaded directly from the GEO database. In total, there are 5 samples from 3 patients, BC09, BC10 and BC11. BC10 has the largest overlap with TCGA-derived CDR3 clusters. For BC10, we selected 1,103 genes with standard deviation ≥1 and performed tSNE analysis on the 4,926 cells using these genes for dimension reduction. This filter is purely for visualization purposes. 2-dimensional scatter plot using tSNE values were generated to visualize the distributions of genes of interest. Based on the locally enriched pattern of 18 clustered cells, we defined a subgroup of 44 cells. For each of the 1,103 genes, we performed Wilcoxon rank sum test between this group and other cells and used Benjamini-Hochberg method to evaluate FDR. These results, including the cell barcodes for the selected group, are available in **Supplementary Table 3**. ZNF683 expression levels in the TCGA samples were split into two groups by the median level. Survival analysis for ZNF683 was performed using Cox proportional hazard model on the binary variable corrected for patient age.

We performed cell trajectory analysis for selected clonotypes in the breast cancer samples. For sample BC10, we selected 418 cells with CDR3 sequences found in the CD8+ subgroup identified in the tSNE plot, and used R package monocle^48^ to perform cell ordering by pseudotime. As the direction of pseudotime is arbitrary, we used representative biomarkers for T cell activation to determine the beginning of the trajectory, and identified the T_pre_ population. The T_rm_ clusters were then selected at the end of the trajectory. Spearman’s correlation between each gene expression level and pseudotime was calculated, and we selected important biomarkers for cell identity (IL7R, TCF7, CCR7), cytotoxicity (GZMB, PRF1, IFNG), exhaustion (PD-1, LAG3, TIM-3), resident memory signature (SELL, KLRG1, CD103) and metabolic status (OXPHOS genes). For BC11, we first merged the two biological replicates into one dataset and selected 31 cells with IL7R≤1, TCF7≤1, GZMB≥5, ZNF683≥5 and CD103≥10 as tissue resident T cells, and used all the 728 cells sharing the same CDR3s with these 31 cells to perform the pseudotime trajectory analysis. These cells in total come from 11 clonotypes, but for the individual clonotype evolution analysis, we removed two clonotypes with n=1 and showed the remaining 9 in **Supplementary Figure 8c**. We did not identify any cell using the same selection criteria for tissue resident memory cells for sample BC09.

### Gene expression analysis

We performed a genome-wide correlative analysis to identify genes associated with counts (*K*) of clustered CDR3s in each individual (**Supplementary Figure 4**). We first selected 15 cancer types with sufficient sample size (≥100). For each cancer, we calculated partial Spearman’s correlation between *K* and the expression level for each gene. Tumor purity is corrected in this analysis as it is expected to impact gene expression profiles^63^ and is correlated to T cell infiltration. False discovery rate is estimated using Benjamini-Hochberg procedure for all the p values pooled.

We also performed differential gene expression analysis to identify novel cancer associated antigens (**Figure 4**). First, we selected 120 clusters with CDR3 length 20≥L≥13 and with ≥10 sequences. For each cluster, we performed one-tailed Wilcoxon rank sum test for each gene between clustered and non-clustered individuals from all cancers, pooled all the p values and estimated FDR using Benjamini-Hochberg correction. This step selected 3,524 significant results (FDR<0.05 and fold change ≥10), including 1,409 unique genes spanning 115 clusters. Fold change was calculated for each cluster, as the median expression value of the samples in the CDR3 cluster divided by that of those not in the cluster. If the denominator is zero, we used an arbitrarily small number 10^-13^. Of all the protein coding genes, HSFX1 has the top significant value, and is associated with clusters 1724 and 1767. We performed a second differential gene expression analysis to visualize the top highly expressed genes, by combining samples in the two clusters.

### Vaccination of naïve and transgenic mice and ELISPOT assay

C57BL/6J and C57BL/6J-HLA-A2.1Tg mice were purchased from the Jackson Laboratory. All mice were maintained under specific pathogen–free conditions at UT Southwestern Medical Center. 10 μg of VMF or VRF peptide was mixed with 50 μg ODN1826 and 100 μg poly (I:C) in 100 μl PBS and then subcutaneously injected to the mouse on day 0 and day 14. Single cell suspensions were prepared on day 18 post first vaccination. Splenocytes were seeded at 4*10^5^ per well and stimulated with either 10 μg peptide or PMA + Ionomycin for 36 hours. ELISPOT assay was performed using an IFN-γ ELISPOT assay kit (BD Biosciences) according to the manufacturer’s instruction. Spots were enumerated by ImmunoSpot Analyzer (CTL).

### Analysis of missense and frameshift mutations

In total we analyzed 920,483 somatic missense mutations from 7,046 TCGA samples with whole exome sequencing data. Mutations occurred in fewer than 3 individuals were excluded, resulting in 6,136 mutations across 5,774 individuals. For each mutation, we estimated its co-occurrence with each of the 671 clusters with CDR3 length 20≥L≥14 and with ≥3 sequences. The length cutoffs were applied to minimize the inclusion of public TCRs, or potentially incorrect CDR3 assemblies. The cluster size cut-off was applied to select those with potentially sufficient statistical power. For each comparison, Fisher’s exact test was performed to estimate a p value (p), only when there is an overlap between the mutation carriers, and those in the CDR3 cluster. It is clear that the p values generated in this approach were inflated due to the selection criterion. Therefore, we conducted a permutation analysis to evaluate the real significance levels and correct for multiple hypothesis testing.

We initialized loop counter n=0 and started iterations. Each time, we randomly sampled one CDR3 cluster (denoted as *C*). Let n_c_ denote the number of individuals in *C*, we randomly sampled n_c_ unique IDs from the pool of 5,774 individuals to replace *C*. We then randomly sampled one mutation, and estimated the co-occurrence between *C* and the mutation carriers. If there was no overlap, the loop started over again without changing n. If an overlap existed, we estimated the p value using Fisher’s exact test, and n became n+1. The loop stopped at n=50,000. P values (P_0_) produced from this analysis were used as null distribution to evaluate the corrected significance levels and FDR. Specifically, for each p, we calculated the significance level p’=(number of P_0_ smaller than p)/50,000. FDR was then estimated using Benjamini-Hochberg procedure on p’.

Similarly, we analyzed 53,491 frameshift indels and kept 1,225 ones occurring for ≥3 times. These mutations distributed across 2,810 individuals. We used the same set of CDR3 clusters to study the co-occurrence patterns between mutation and CDR3s, and used the same permutation strategy to evaluate statistical significance and FDR. If one CDR3 cluster was associated with more than one passed-FDR mutations (including indels), we used the most significant one in our downstream analysis. For both types of variants, we selected mutations with FDR≤0.05 and odds ratio from the Fisher’s exact test ≥1000. The cut-off in the odds ratio was applied to select highly specific enrichment of mutations in the CDR3 clustered individuals.

### HLA allele binding prediction

All the HLA allele binding predictions in this work were performed using either NetMHC or NetMHCpan online server. We implemented NetMHCpan for less common HLA alleles not covered in NetMHC. For missense mutations, the input peptide is a 17-mer with mutated amino acid in the middle. For frameshift mutations, we included 8-mer before and all the amino acid sequence after the mutation locus. For cancer-associated antigens, we downloaded the complete protein sequence from Uniprot (www.uniprot.org), and input the fasta file to NetMHC/NetMHCpan server. Biding of the control peptide VRF for *in vivo* validation to HLA-A*02:01 was predicted using NetMHC server. Default rank cut-offs were applied to define weak (≤2) or strong binders (≤0.5).

### Prediction of cancer disease status

In this analysis, we compared 3 TCR repertoire datasets from different studies, including pre/post anti-CTLA4 treatment late stage melanoma (melanoma), early breast cancer (breast cancer), and HCMV cohort (HCMV) as normal control. To avoid systematic bias after public TCR removal, we used second batch (n=120) of the HCMV cohort in this analysis. Direct comparison between different study cohorts will be biased towards sequencing depth and the amount of lymphocytes captured for sequencing. Therefore, we conducted a downsampling procedure to ensure the comparability across cohorts. The targeted capture protocol applied for TCR repertoire sequencing allowed one read to completely cover the whole CDR3 region. Therefore, read count is used to estimate clonal abundance from the Adaptive Biotechnology immunoSEQ Analyzer. We first calculate the size for each TCR-seq library (N), which is the summation of the read counts (*m*) for all the CDR3 calls. A combined vector of CDR3s with length N was made, with each CDR3 sequence *i* repeated by *m_i_* times, where *m_i_* is the read count for CDR3 sequence *i*. For the melanoma cohort, we used all the cancer samples (n=21 for either pre- or post- treatment), and randomly sampled 100 individuals with replacement as normal control. For each of the 121 samples, we downsampled the library to K=100,000 reads, each read being a CDR3 amino acid sequence. The read count (*m*′) for each unique CDR3 was then calculated. For each sample, CDR3s with identical sequence to one of the cancer-associated CDR3s were selected, and the summation of *m*’ for all the selected CDR3s was used as predictor for cancer status. For breast cancer cohort, same downsampling strategy was applied, except that we used K=30,000 for PBMC and 10,000 for TIL samples.

### Statistical Analysis

Statistical analyses were performed using R statistical programming language^64^. Survival analysis was implemented using Cox proportional hazard model in R package *survival*. Receiver operator curves and area under curve calculations were performed with R package *AUC*. tSNE plots for single cell analysis were generated using *Rtsne*^65^. Single cell pseudotime trajectory analysis was performed using *cellrangerRkit* and *monocle*^48^. The statistical significance for Figure 5c was estimated separately. The genotype frequency for A*02:01 in the TCGA cohort is 0.417, and the frequencies for C*07:01 and C*07:02 are smaller. We used A*02:01 frequency to estimate a conservative p value: the probability of observing 9 individuals carrying A*02:01 is 0.417^9^=0.00038. Therefore, the p value for observing the configuration in Figure 5c is significant. Similarly, the most abundant allele type in **Supplementary Figure 13c** is C*07:01, with genotype frequency 0.253, and the p value for this configuration is ≤ 0.253^5^=0.0010. Two-way ANOVA test for comparing different treatment groups of vaccinated mice was performed using commercial software GraphPad Prism.

**Supplementary Figure 1.**
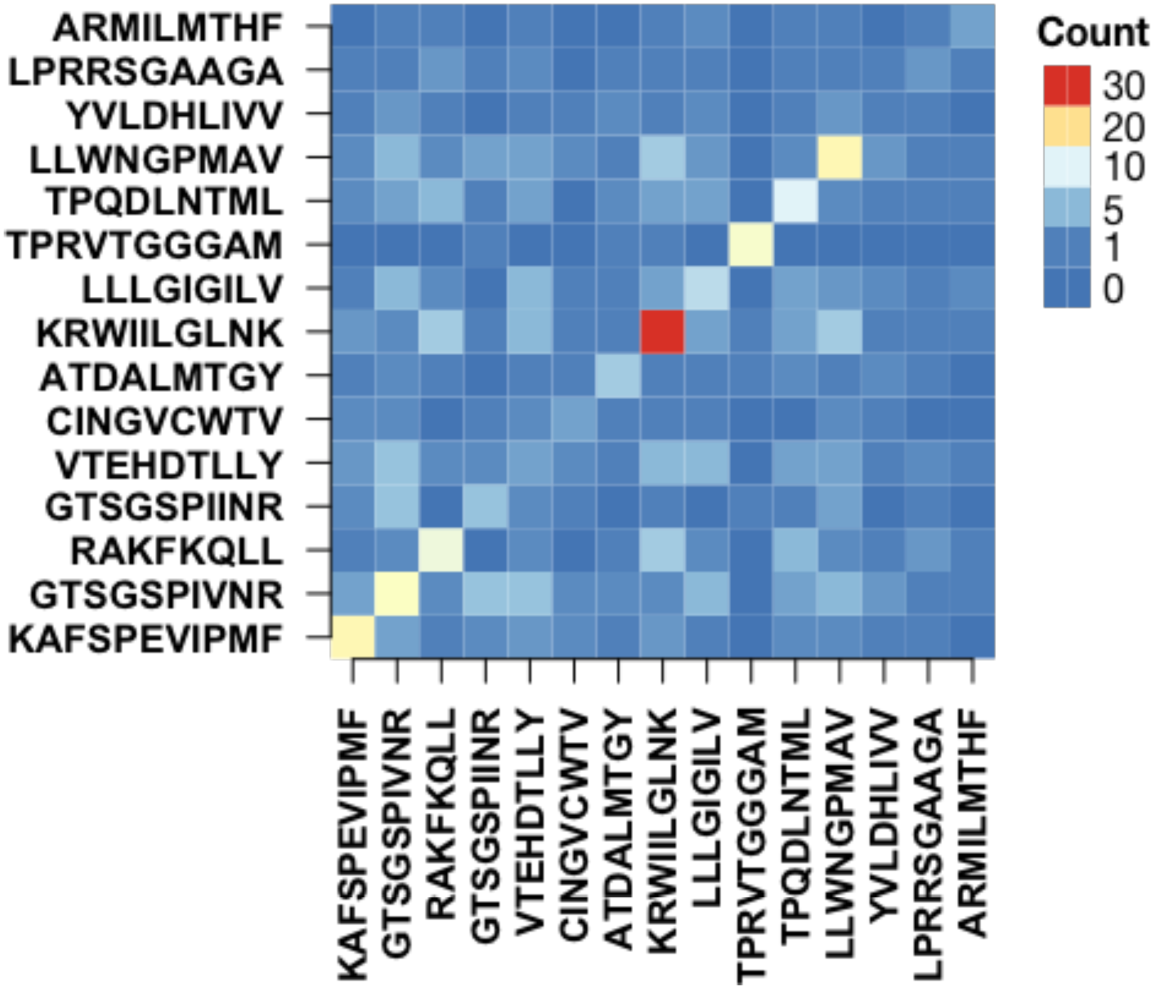
Cluster antigen-specificity analysis for GLIPH. Heatmap showing the cross-antigen classification errors from GLIPH predicted CDR3 clusters using the same benchmark dataset of 15 antigens. Same analysis was performed as described in **Figure 1b**.

**Supplementary Figure 2.**
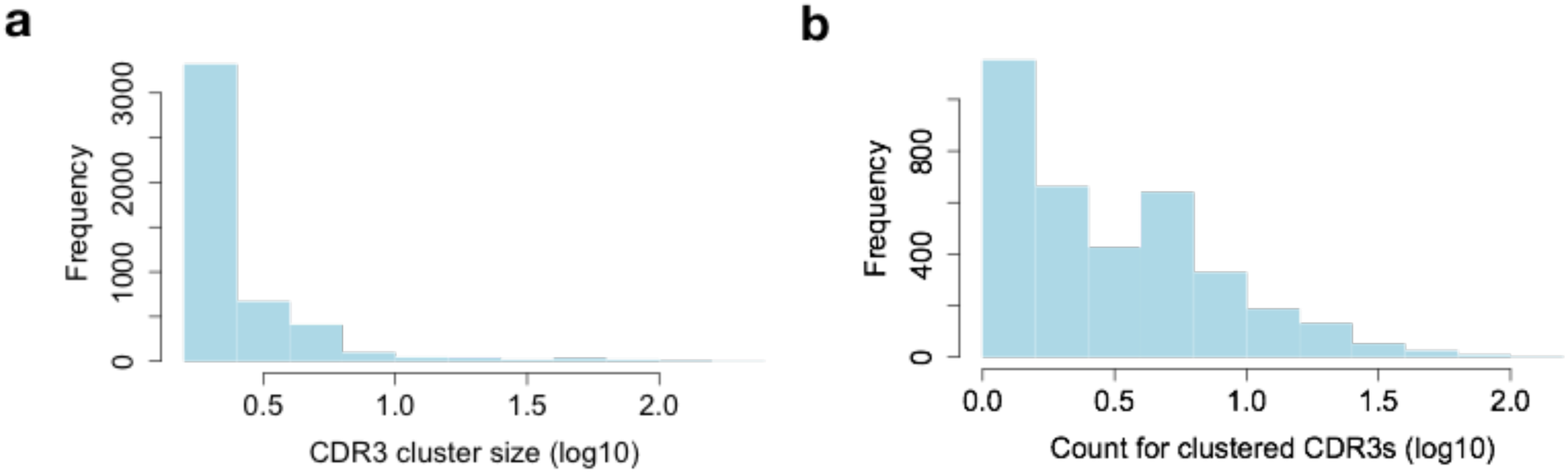
Summary of iSMART identified CDR3 clusters. **a**) Histogram of CDR3 cluster size distribution. **b**) Lengths distributions for clustered and non-clustered CDR3 amino acid sequences. **c**) Distribution of the counts for clustered CDR3s carried by each individual in the analysis.

**Supplementary Figure 3.**
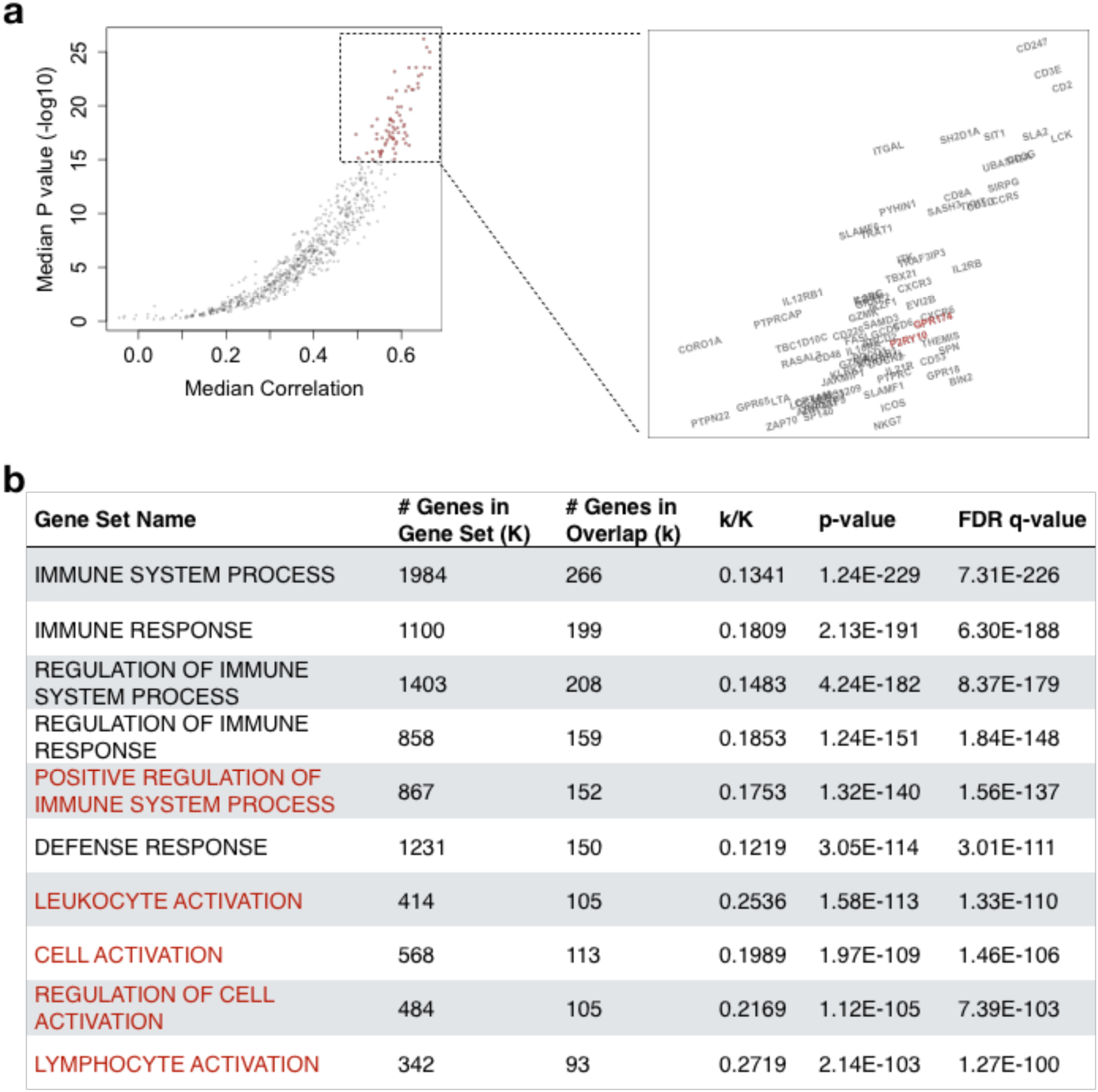
Clustered CDR3s as an indicator for activated T cells. **a**) Volcano plot for genes positively correlated with number of clustered CDR3s. Median values across different cancer types for each gene were calculated for both p value and partial Spearman’s correlation with tumor purity correction. Top genes (p≤10^-15^) were zoomed in for visualization. Genes related to negative regulation for Treg cells were highlighted with dark red color. **b**) Gene-Ontology enrichment analysis was performed for the top 500 genes and the top 10 pathways were displayed. Highlighted pathways were related to immune cell activation.

**Supplementary Figure 4.**
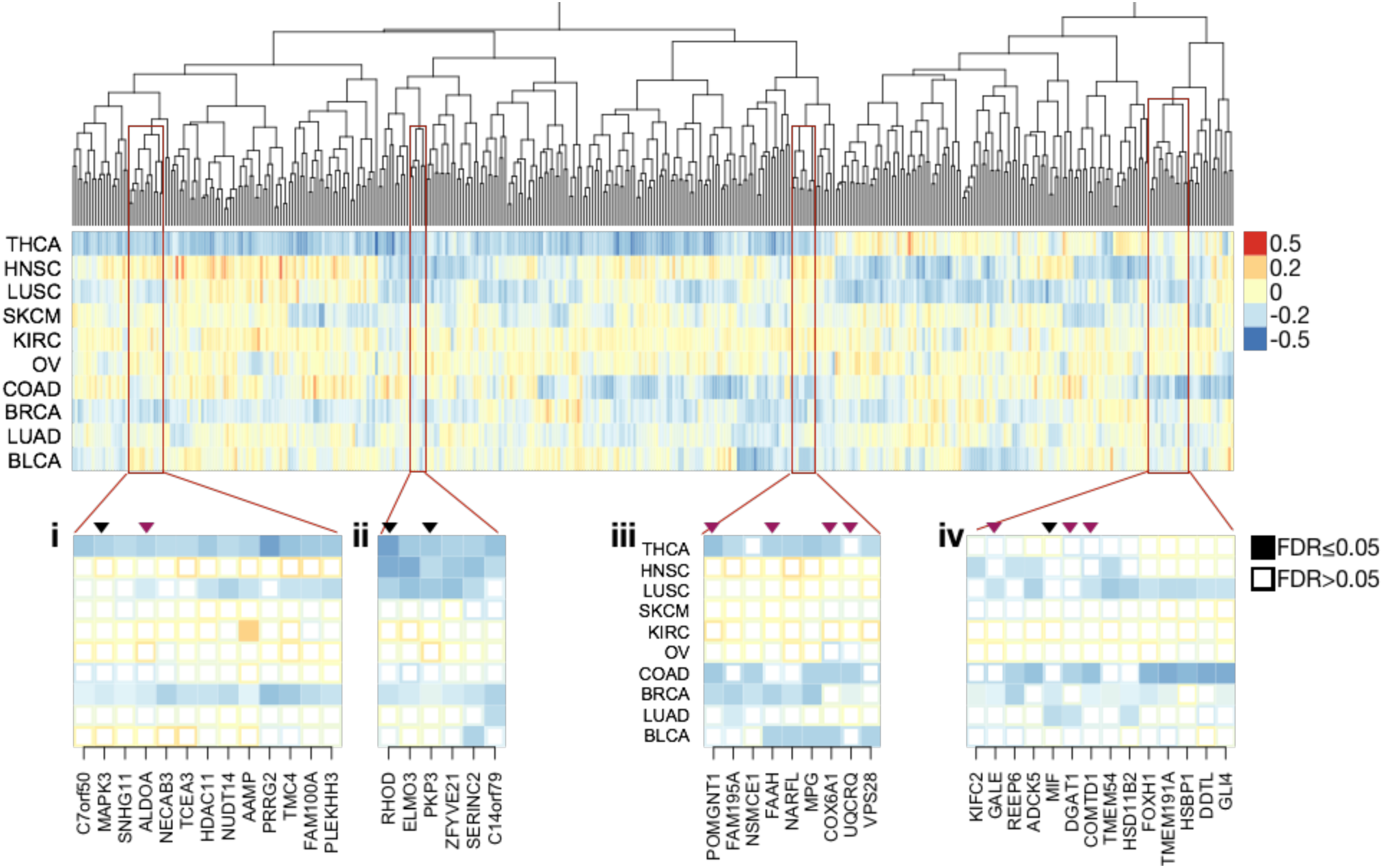
Potential negative regulators for T cell activation in the tumor microenvironment. Genes with Spearman’s correlation ρ≤ -0.1 and FDR≤0.05 in at least 3 cancer types were selected for visualization in the heatmap. Hierarchical clustering on ρ was performed to order the genes into similar groups across different cancer types. Four representative clusters with putative oncogenes (labeled by black arrows) or recently identified metabolic enzymes (red arrows) were displayed as smaller heatmaps in the lower panels. Statistical significance was evaluated using partial Speaman’s correlation test correcting for tumor purity, and FDR was performed using Benjamini-Hochberg procedure.

**Supplementary Figure 5.**
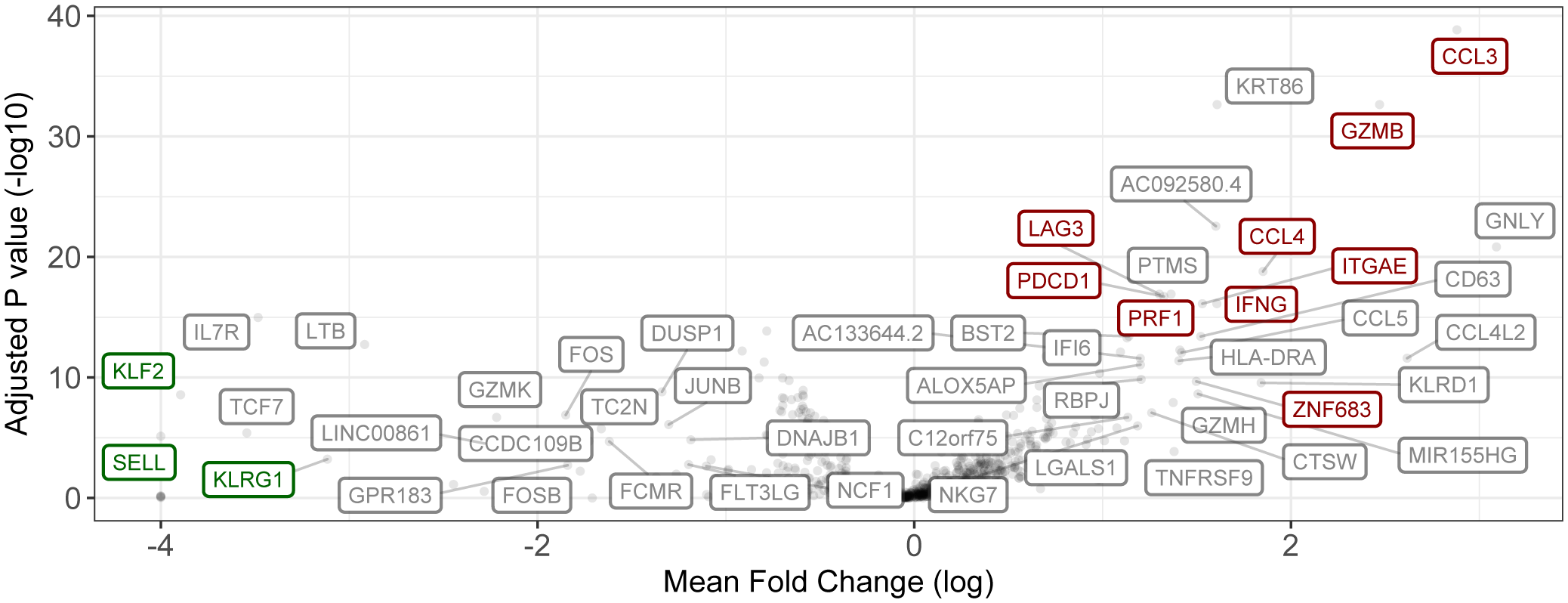
Differentially expressed genes between our defined new group and other T cells. Wilcoxon rank sum test was applied to evaluate the statistical significance, and p values were corrected by Bejamini-Hochberg method. We labeled the genes with mean fold change greater than 3 or smaller than -3, and FDR<=0.01. Established markers for tissue-resident memory T cells were highlighted with colors: red for up-regulation and green for down-regulation.

**Supplementary Figure 6.**
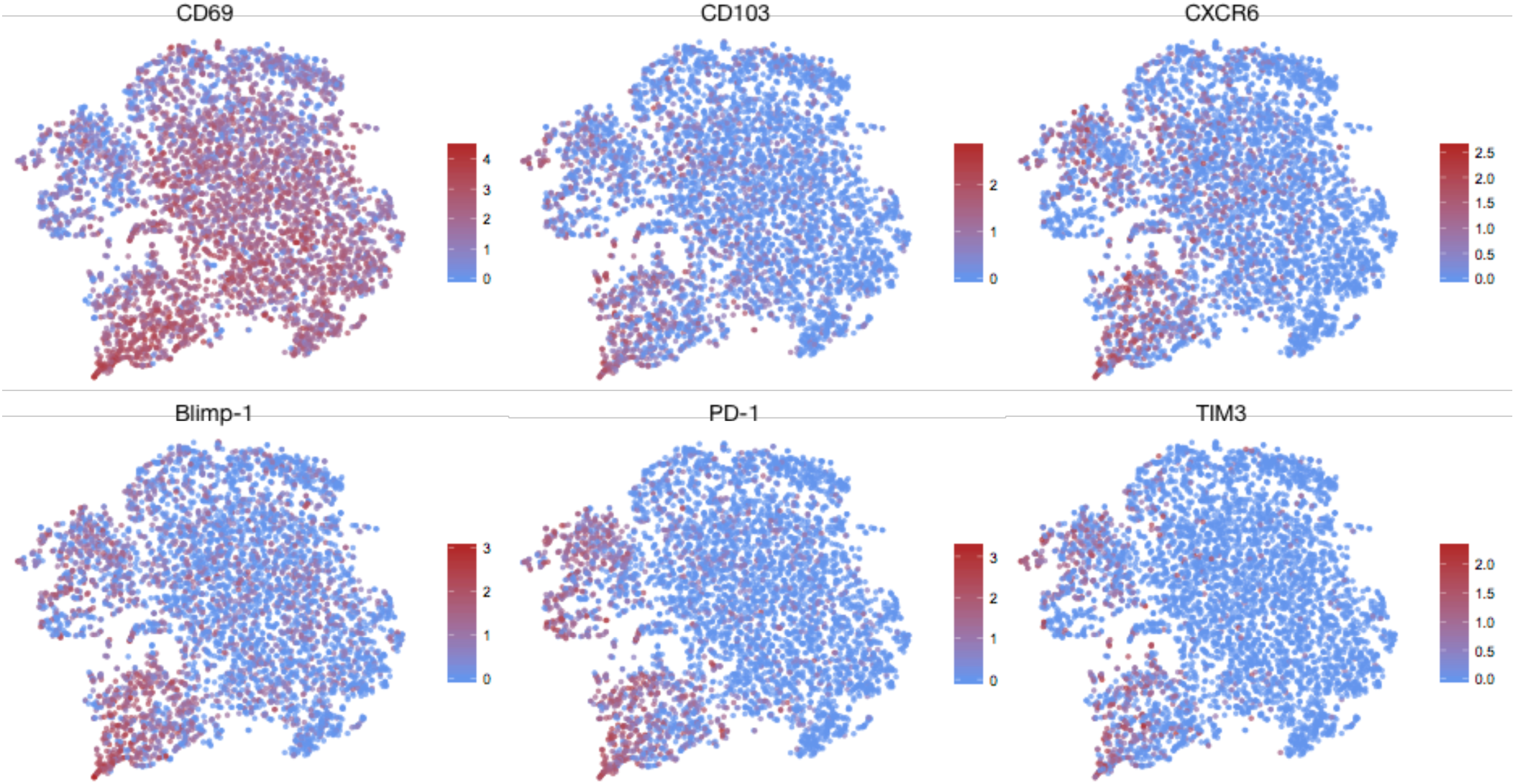
Additional tSNE plot visualization for selected markers. Previously reported Trm markers including CD69 (general T cell activation and memory differentiation), CXCR6, Blimp-1 and CD103 (ITGAE) were visualized by tSNE plots. Expression patterns for two putative T cell exhaustion markers, PD-1 and TIM3 were also presented. Figure legend was in log scale.

**Supplementary Figure 7.**
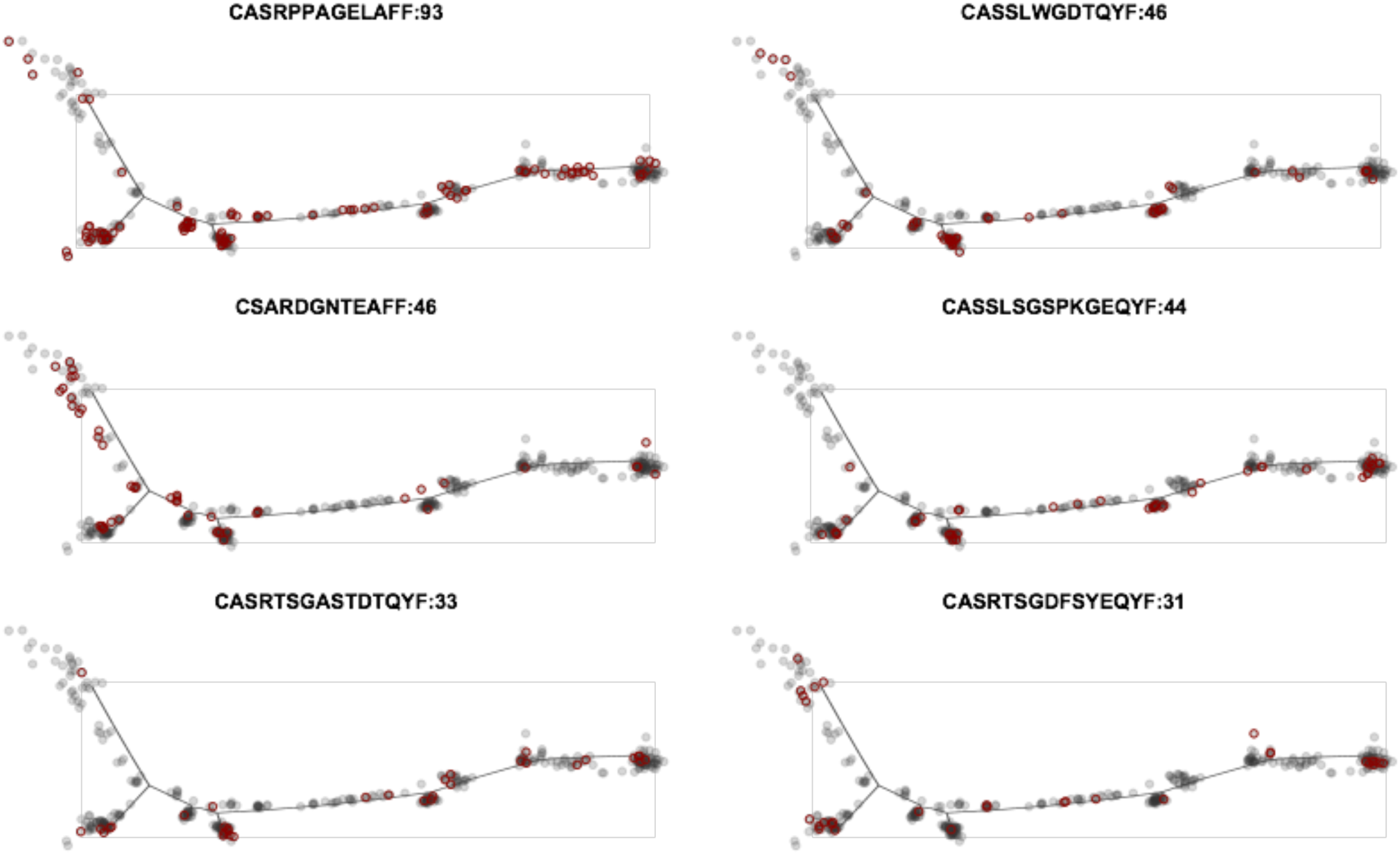
Pseudotime trajectory plots for individual clonotypes in sample BC10. βCDR3 sequences were displayed as figure titles and each point on the plot represent a cell. Red circles label cells with the corresponding CDR3 sequence. The numbers in the figure titles are the number of cells in the corresponding clonotype. Two evolutionary patterns were observed. Pattern 1 is from T_pre_ to T_rm1_, including clonotypes CASRPPAGELAFF, CASSLWGDTQYF, CASSLSGSPKGEQYF, CASRTSGASTDTQYF and CASRTSGDFSYEQYF. Pattern 2 is from T_rm1_ to T_rm2_, including clonotype CSARDGNTEAFF.

**Supplementary Figure 8.**
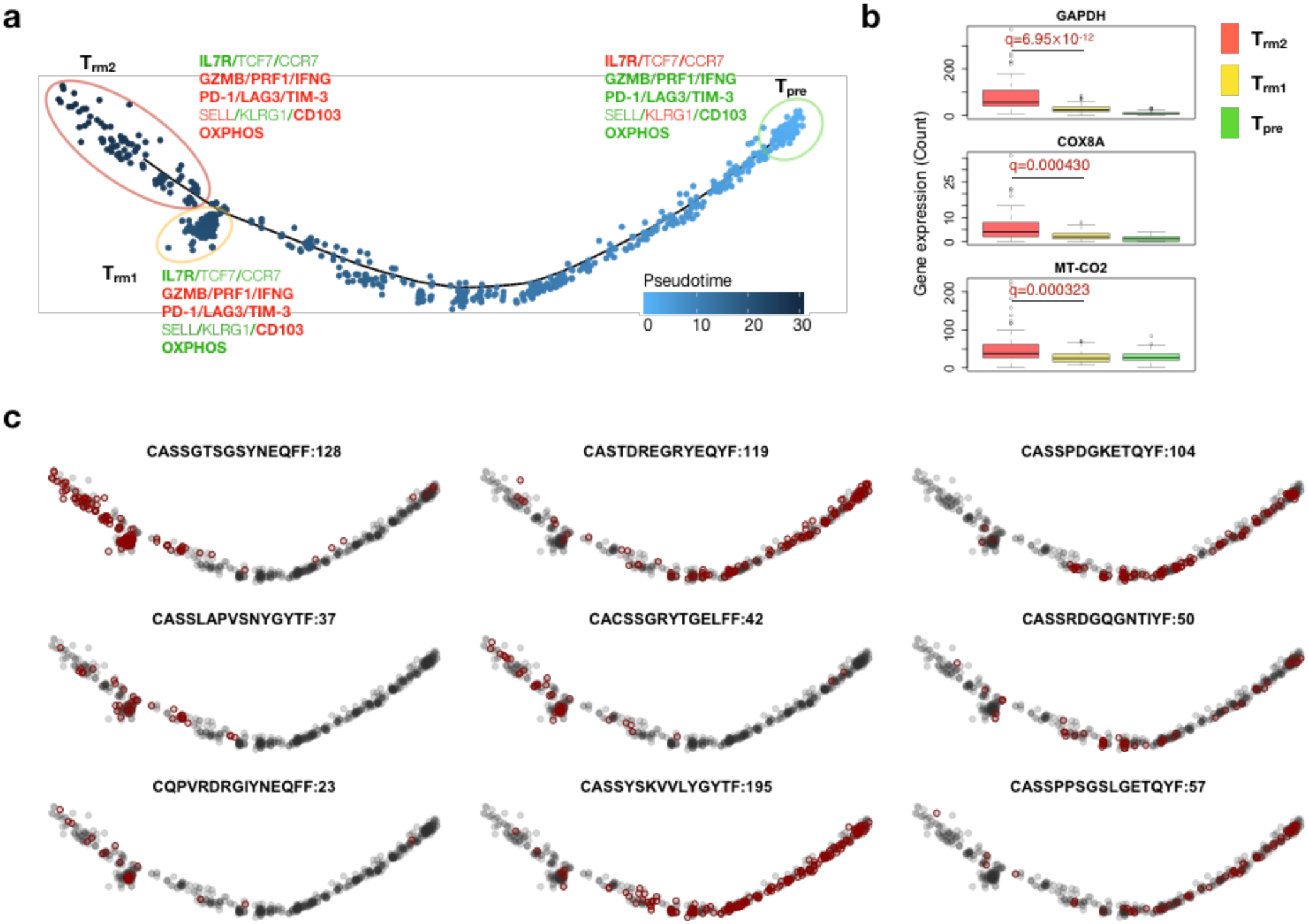
Single cell trajectory analysis for breast cancer sample BC11. **a**) Same trajectory analysis in BC11, with exception that for the selected markers of the cell clusters, bold characters indicate FDR<0.05, where normal font otherwise. **b**) Boxplot for representative OXPHOS genes, with statistical significance evaluated by Wilcoxon rank sum test. **c**) Trajectory plots with individual clonotype overlaid by red circles. The number after each CDR3 sequence is the number of cells in the corresponding clonotype. Two evolutionary patterns were observed. Pattern 1 is from T_pre_ to T_rm1_, including clonotypes CASTDREGRYEQYF, CASSPDGKETQYF, CASSRDGQGNTIYF, CASSYSKVVLYGYTF and CASSPPSGSLGETQYF. Pattern 2 is from T_rm1_ to T_rm2_, including clonotypes CASSGTSGSYNEQFF, CASSLAPVSNYGYTF, CACSSGRYTGELFF and CQPVRDRGIYNEQFF.

**Supplementary Figure 9.**
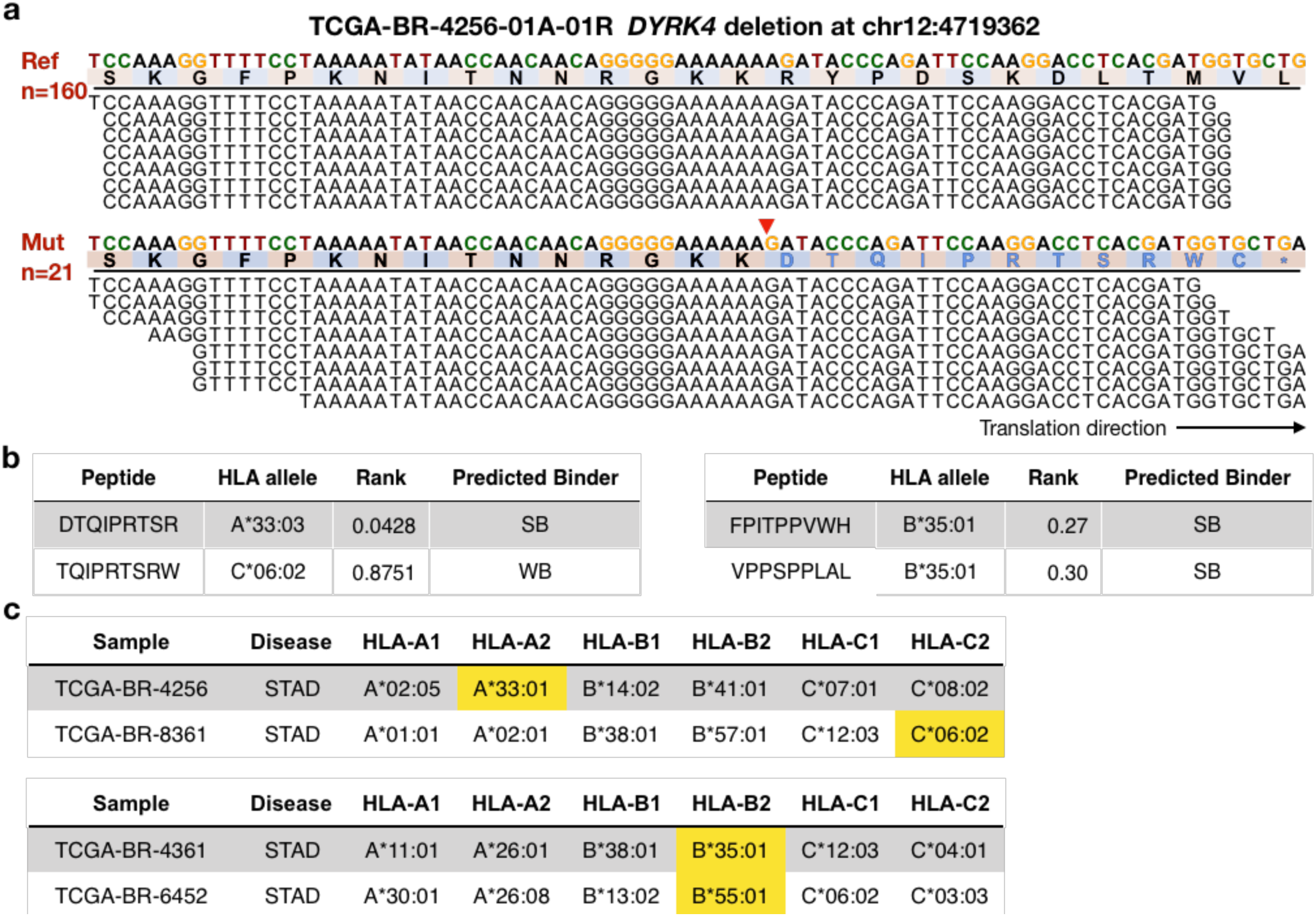
Potential neoantigens generated from other top frameshift indels. **a**) Read pileup plot for gene *DYRK4* frameshift deletion. The gene is forward translated. Site of deletion was labeled with red arrow. **b-c**) NetMHC predictions for binding peptides and TCGA sample HLA genotypes: left table in **b** and upper table in **c** for gene *DYRK4*; right table in **b** and lower table in **c** for gene *RNF43* in **Figure 3c**.

**Supplementary Figure 10.**
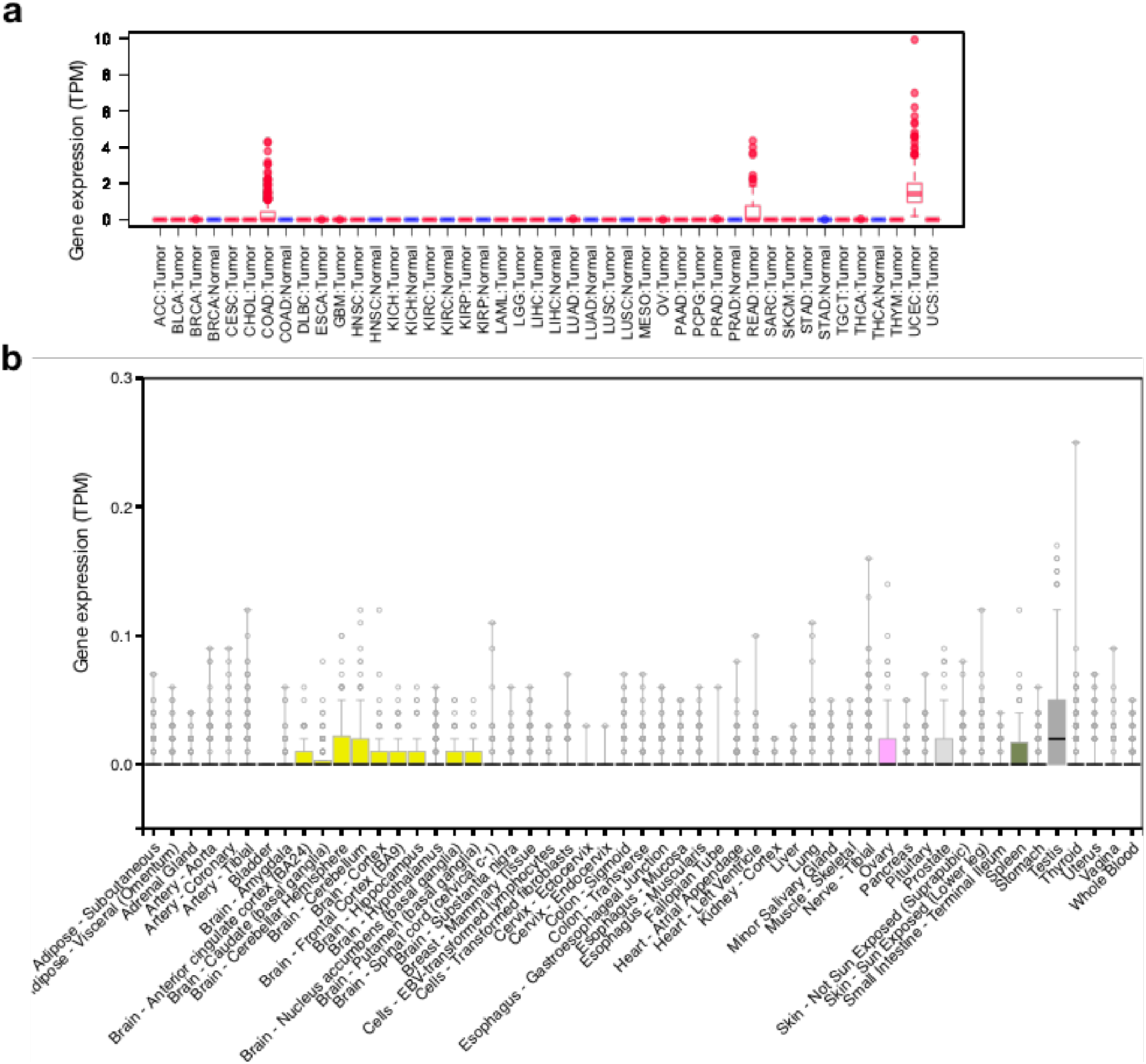
*HSFX1* expression across cancer types and normal tissues. **a**) TPM values for HSFX1 expression were displayed in boxplots across 32 cancer types. Adjacent normal samples with sufficient sample size (n≥20) were also included in the plot. **b**) TPM values for HSFX1 expression displayed in boxplot across 53 normal tissue types as reported by the GTEx data portal. Box colors distinguish major tissue types: yellow for brain, pink for ovary, light gray for prostate, dark gray for testis and olive green for spleen.

**Supplementary Figure 11.**
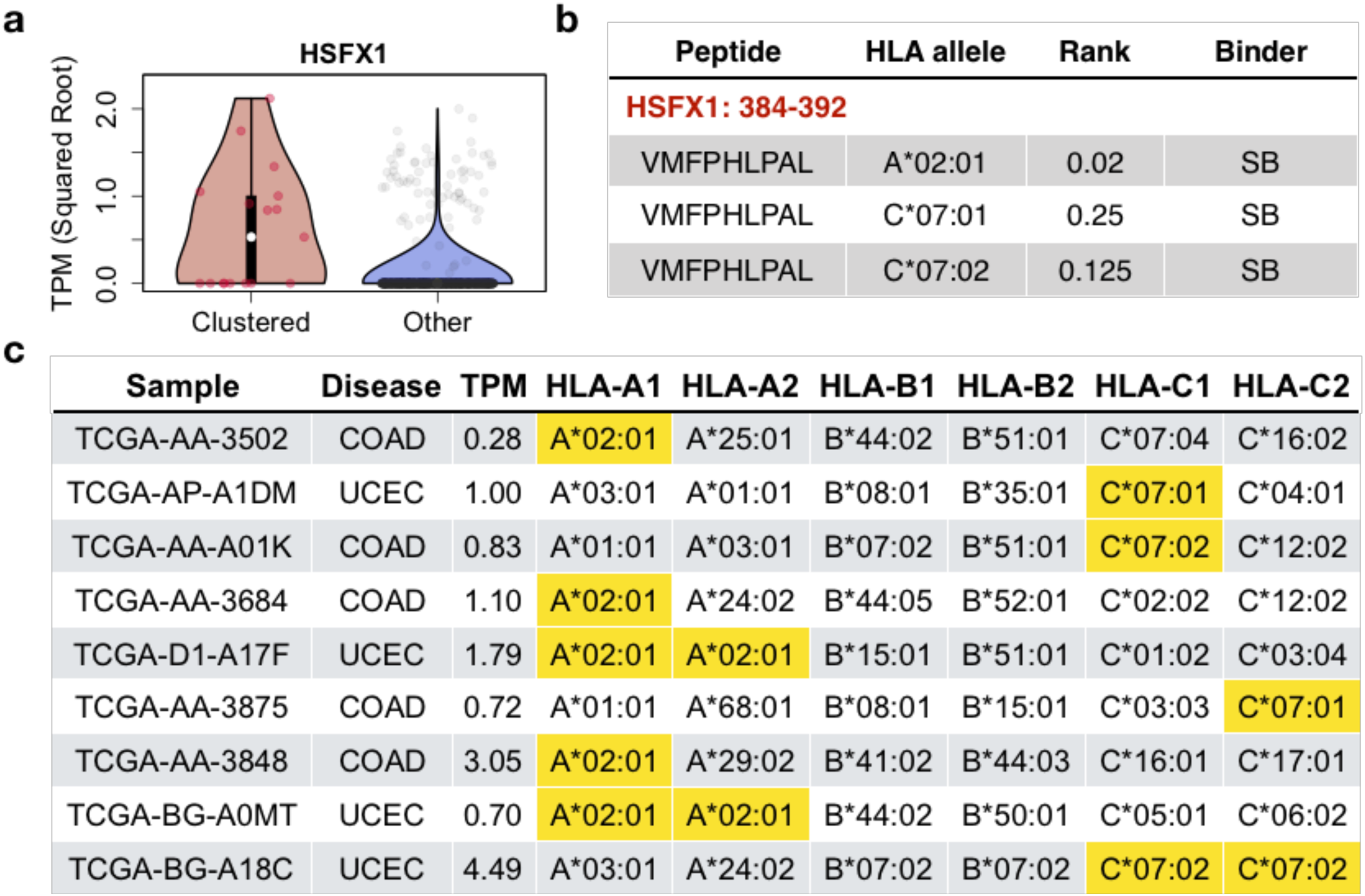
Generation of a potentially immunogenic peptide by *HSFX1* expression. **a**) Violin plot showing significant difference in *HSFX1* expression between clustered and non-clustered individuals from all TCGA cancers. 9 individuals with solved HLA genotypes have positive *HSFX1* expression. **b**) NetMHC prediction for a *HSFX1* derived 9-mer peptide, with strong binding affinities to 3 common HLA alleles. **c**) HLA genotype information table for the 9 individuals with *HSFX1* expression shown in **a**). All 9 samples were colon or endometrial cancers, bearing at least one matched HLA binder(s) as shown in **b**) (highlighted in yellow). Expression levels in transcript per million (TPM) were also displayed in the table.

**Supplementary Figure 12.**
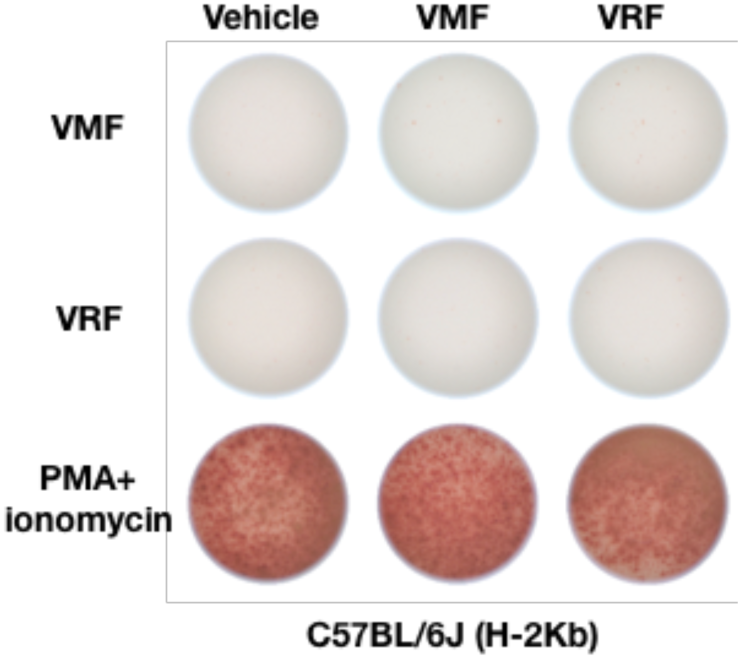
Control experiment for *HSFX1*-derived peptide using naïve C57BL/6J mice. Same experiment as described in Figure 6a was performed using naïve immunocompetent C57BL/6J mice (n=4, all female), and representative ELISPOT results were displayed.

**Supplementary Figure 13.**
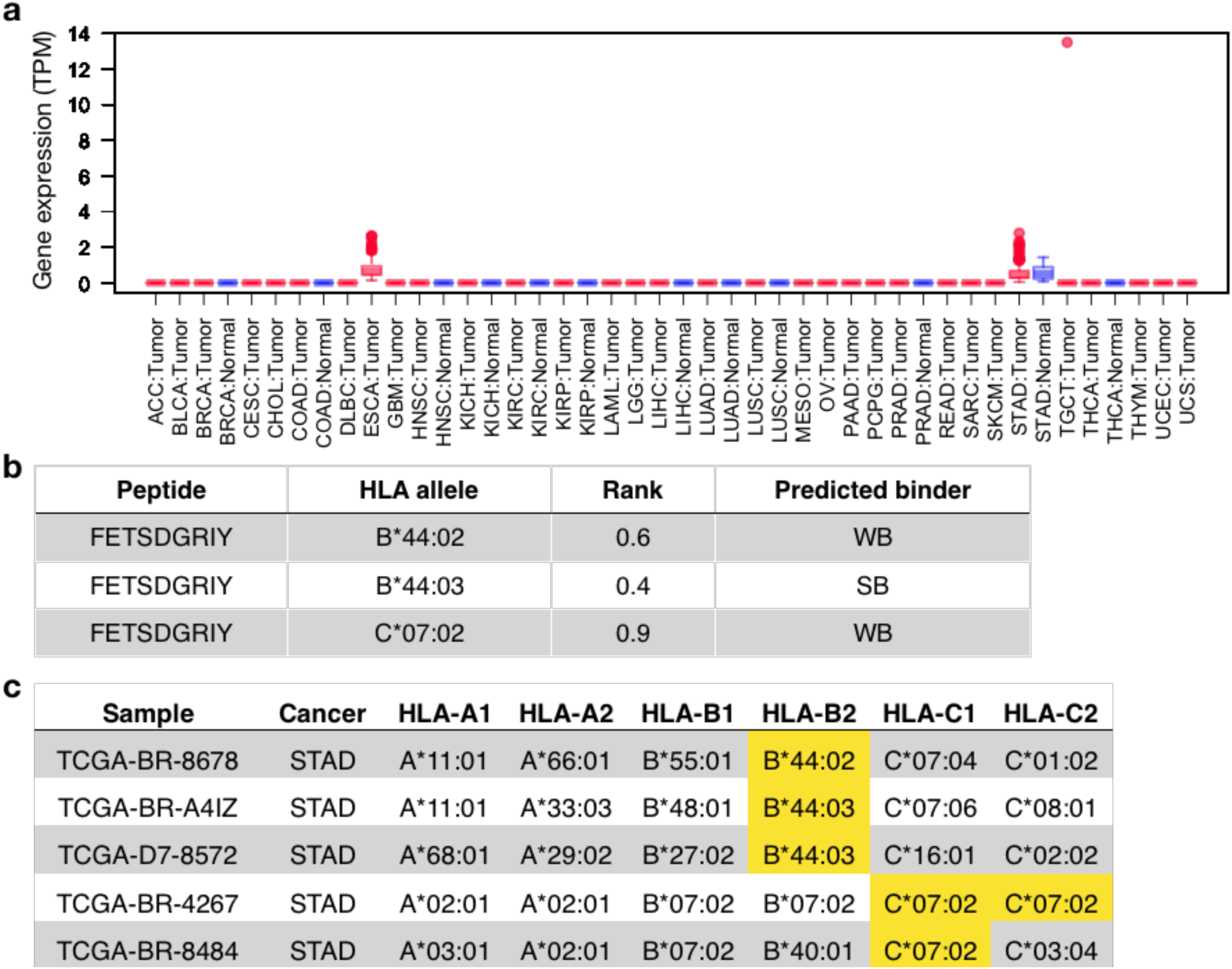
Identification of *TSSK2* as a potential cancer-associated antigen. **a**) Boxplot for *TSSK2* expression across different tumor and normal tissues. **b-c**) NetMHC predicted binding peptide information and HLA genotypes for samples with *TSSK2* expression.

**Supplementary Figure 14.**
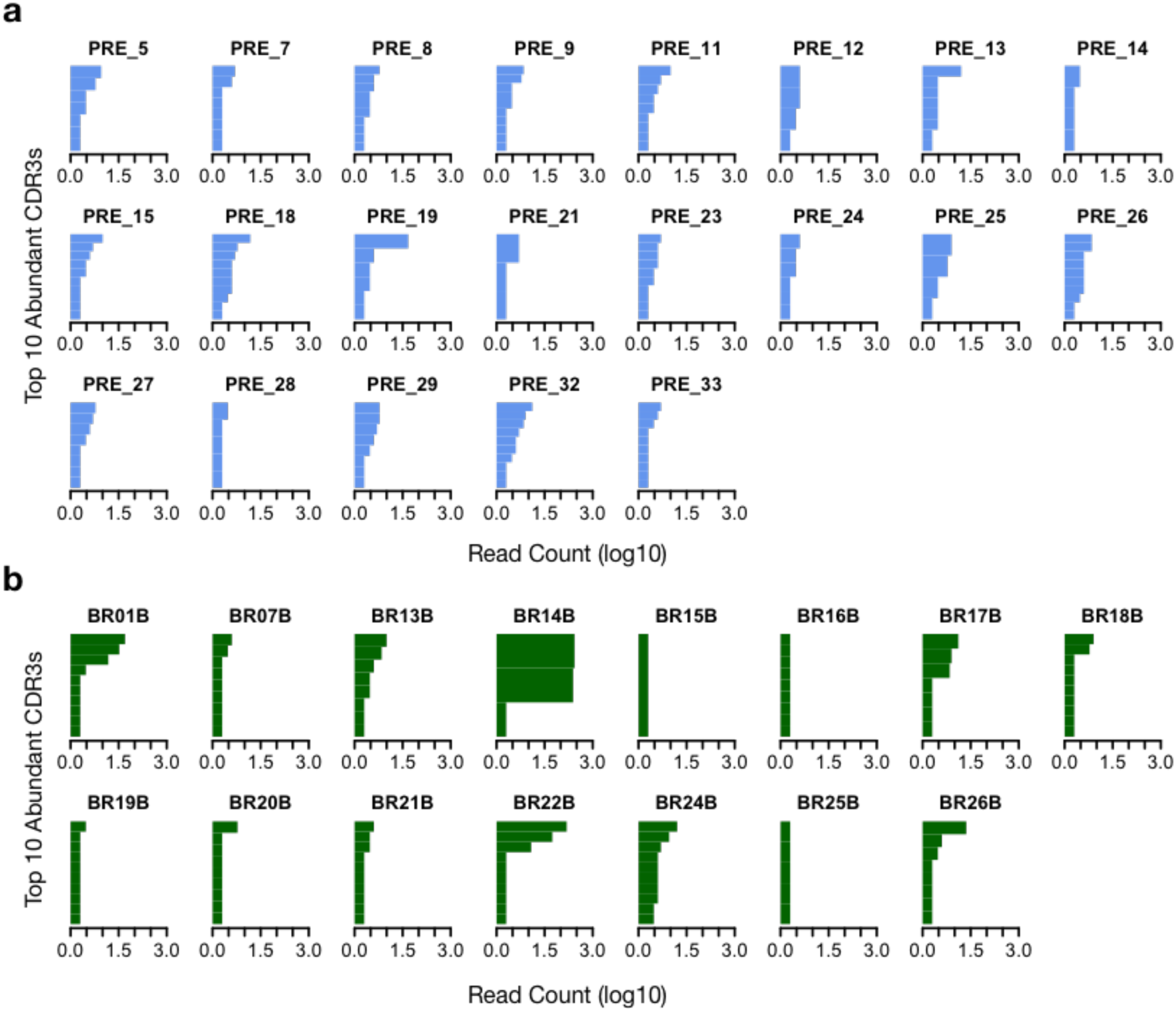
Cancer-associated T cell clonal frequencies in early and late stage tumors. **a-b**) Barplots showing the read counts for top 10 cancer-associated CDR3s in samples of the melanoma (**a**) or early breast cancer (**b**) cohorts. Since the library sizes for TCR-seq data in the melanoma cohort were bigger than those for the early breast cancer cohort, to ensure fair comparison, we downsampled the libraries of the melanoma samples to *N,* which randomly sampled from Uniform distribution (34383,156524). The upper and lower limits mark the range of library sizes for early breast cancer samples. Specifically, for each library, we randomly sampled an integer (N) from that range, and downsampled the initial library to N reads, and estimated CDR3 frequencies (**Methods**). The above range matches the library sizes for the early breast cancer cohort.

**Supplementary Table 1.** Information for the 15 selected antigens for methodology performance evaluation.

**Supplementary Table 2.** Summary of association analysis between gene expression levels and clustered CDR3 count in each individual.

**Supplementary Table 3.** Summary of differential gene expression analysis between the single cells from new defined group and others, with fold change and FDR estimations.

**Supplementary Table 4.** HLA genotype information for patients carrying correlated pairs of predicted neoantigens and clustered CDR3s.

**Supplementary Table 5.** Summary of differential gene expression analysis between each CDR3 cluster and other TCGA tumors, with fold change and FDR estimations.

**Supplementary Dataset 1.** TCR clusters obtained from the top 5000 most abundant clonotypes of 666 HCMV cohort. (Due to file size limit, in this submission we only included 3 files for review. Format of the remaining files is the same.)

**Supplementary Dataset 2.** Public non-specific TCRs profiled using random triplets with unmatched HLA alleles from the HCMV cohort.

